# Bifunctional methacryloyl-norbornene gelatin chemistry enables tunable bioinks for soft tissue engineering via digital light processing

**DOI:** 10.64898/2026.09.17.752269

**Authors:** I. Pokholenko, A. Lapere, R. Willemarck, N. Eeckman, L. Van Damme, I. Dehaene, S. Van Vlierberghe, N. Pien

**Affiliations:** Polymer Chemistry and Biomaterials Group, Centre of Macromolecular Chemistry (CMaC), Ghent University, Ghent, Belgium; Department of Cell Regulatory Mechanisms, Institute of Molecular Biology and Genetics of NASU, Kyiv, Ukraine; 4Tissue, Zwijnaarde, Belgium; Department of Obstetrics and Gynaecology, Ghent University Hospital, Ghent, Belgium

**Author notes:** Corresponding author: Dr. Nele Pien.

**Keywords:** Digital light processing (DLP), Gelatin, Thiol-ene crosslinking, Bioink, Soft tissue engineering

## Abstract

Photocrosslinkable gelatin derivatives are promising bioinks for soft tissue engineering, but their use in cell-laden digital light processing (DLP) requires a balance between tunable mechanics, printability and cytocompatibility. Here, we report a small library of photocrosslinkable gelatin-based precursor formulations for cell-laden DLP, with the broader aim of generating a versatile platform for soft tissue engineering rather than a formulation restricted to one tissue model. A bifunctional methacryloyl-norbornene gelatin (GelMANB) was developed to combine methacryloyl chain-growth polymerisation with norbornene-mediated thiol-ene crosslinking and was benchmarked against gelatin methacryloyl (GelMA). GelMANB formulations containing different thiolated crosslinkers were evaluated to determine the influence of crosslinking chemistry as well as polymer concentration on network formation and physico-chemical properties. Thiol-ene formulations enabled broad tuning of hydrogel stiffness and exhibited rapid photocrosslinking, while crosslinker selection further influenced swelling and tensile behaviour. Selected GelMANB/dithiothreitol (GelMANB/DTT) and GelMANB/thiolated gelatin (GelMANB/GelSH) formulations were subsequently processed via DLP alongside GelMA. Optimised exposure conditions enabled reproducible fabrication of porous scaffolds that retained dimensional stability under osmotic pressures relevant to several soft tissues. Following cell-laden printing, human foreskin fibroblasts remained viable after seven days, reaching 95.0±5.9% viability in GelMANB/DTT and 94.9±2.7% in GelMANB/GelSH, compared with 87.2±6.0% in GelMA. In 2-mm-thick hydrogel discs, GelMANB/GelSH supported the most sustained fibroblast elongation, reaching 30.9±6.3% elongated cells at day 7, although elongated cells were mainly confined to the outer surface layers. Porous scaffolds with 400 µm struts and 1.3 mm pores reduced the characteristic hydrogel thickness and enhanced elongated morphology across all formulations, highlighting GelMANB as a modular platform for cell-laden soft tissue biofabrication.

**Graphical abstract:** 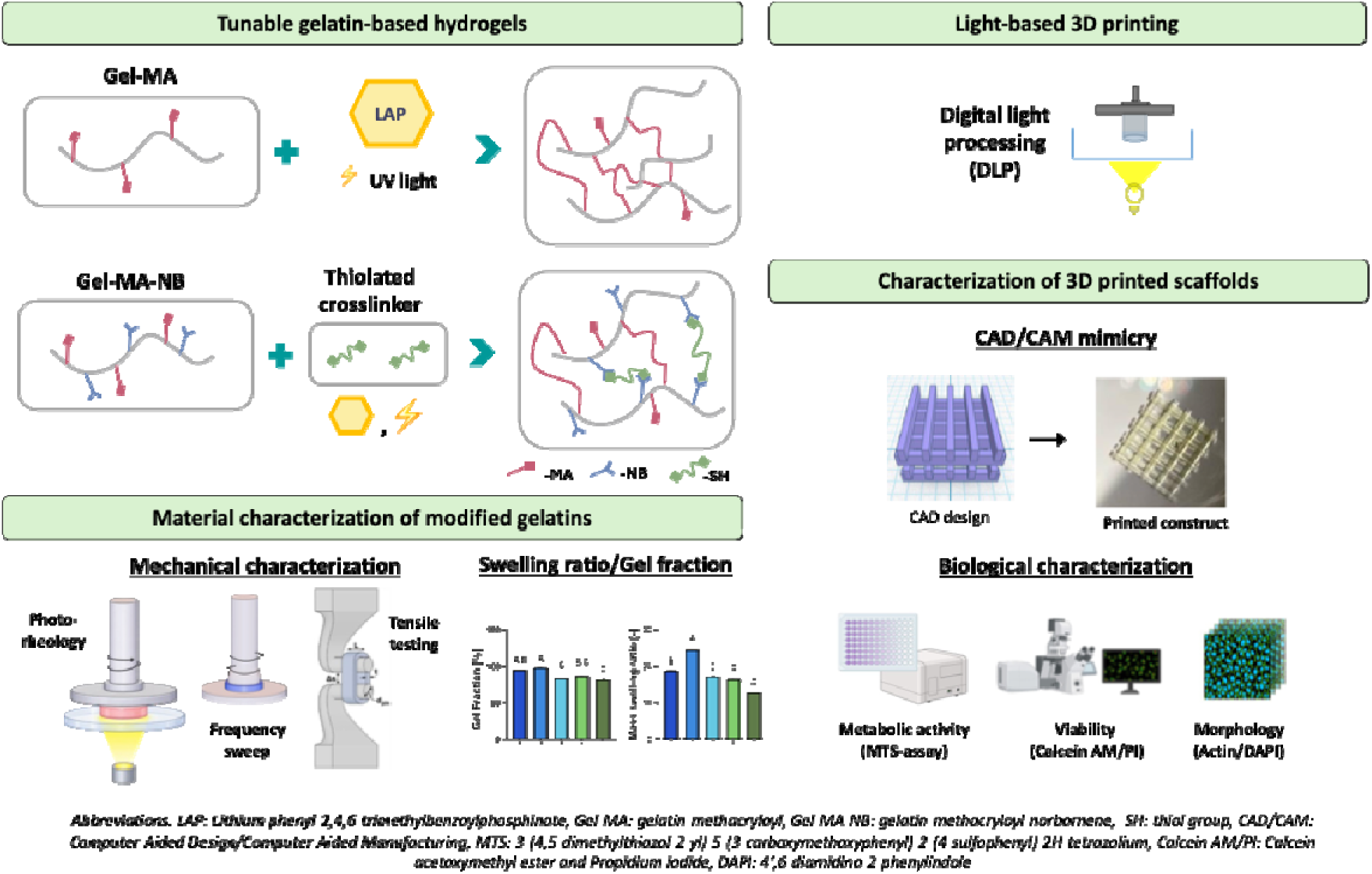

**Highlights:**

- Bifunctional GelMANB enables crosslinker-dependent tuning of hydrogel stiffness, swelling and tensile behaviour.
- GelMANB formulations provide complementary network properties to GelMA while remaining compatible with digital light processing.
- GelMANB/DTT and GelMANB/GelSH improve day-7 fibroblast viability compared with the GelMA benchmark.
- GelMANB/GelSH better support characteristic elongated morphology of fibroblasts than GelMA in cell-laden hydrogels.
- Porous scaffold architecture enhances fibroblast elongation across GelMA and GelMANB formulations by reducing diffusion limitations.

## 1. Introduction

Soft tissues are characterized by high water content, low-to-intermediate stiffness, hierarchical extracellular matrix (ECM) organization and strong cell-matrix reciprocity. These features are essential for tissues that undergo repeated deformation, including lung [1–3], urethra [4–6], vascular [7–9], skin [10,11], adipose [12,13] and smooth muscle-containing tissues [14,15]. In such environments, cells are not only supported by the surrounding matrix, but also actively sense and remodel it through biochemical adhesion motifs, biodegradable sequences and local mechanical cues. As a result, the development of bioinks for **soft tissue engineering** (TE) requires careful balancing of several requirements, including cytocompatibility, cell-interactive chemistry, tunable viscoelastic and swelling behavior, sufficient structural stability after printing and compatibility with (light-based) biofabrication [16,17].

Lung- and urethra-inspired soft tissue engineering provide useful examples of these design requirements, as they represent distinct soft tissue environments with different mechanical, architectural and transport-related constraints. For **lung tissue**, mechanical cues are especially relevant because epithelial cells are continuously exposed to cyclic stretch, compression, shear stress and matrix stiffness changes during breathing, which regulate barrier function, remodeling and repair [18,19]. Human fetal lung elasticity measured using two-dimensional shear wave elastography was reported to range between 3.94 and 5.03 kPa across 24 to 39 weeks of gestation [20]. In this study, the measured region of interest could include both vasculo-bronchial elements and foetal pulmonary parenchyma at earlier gestational ages, whereas measurements after 32 weeks were suggested to more closely reflect the developing pulmonary parenchyma [20]. In adult human lung tissue, atomic force microscopy measurements showed an anatomical stiffness gradient, with elastic moduli of 1.87±0.95 kPa for parenchymal regions, 7.17±4.03 kPa for pulmonary vessels and 15.76±8.88 kPa for airways [21]. Human **urethral tissue** represents a related but distinct soft tissue environment. Cunnane et al. [6] characterized nine human male urethral samples using pressure-diameter and uniaxial extension testing, showing pressure-stiffening, viscoelastic behavior and a correlation between collagen/elastin content and tissue mechanics. Rather than defining urethra by a single modulus value, these data indicate that urethral biomaterials should reproduce nonlinear, viscoelastic, and pressure-responsive mechanical behavior. Together, the human lung and urethral data support the need for precursor formulations that can be tuned across the low-kPa to tens-of-kPa range while maintaining cytocompatibility and processability.

**Hydrogels** are particularly attractive for addressing these requirements because their hydrated polymer networks can reproduce several features of native soft ECMs, including high water uptake, diffusive transport of oxygen and nutrients, and tissue-like mechanical properties [17]. Their composition, crosslinking density and degradation profile can be adjusted to modulate swelling, stiffness, viscoelasticity and cell-matrix interactions, rendering them widely used as precursor systems for soft TE and bioprinting. Among hydrogel-forming polymers, **gelatin** has become one of the most widely explored materials for biofabrication because it is derived from collagen and retains cell-adhesive arginine-glycine-aspartic acid motifs (RGD) as well as matrix metalloproteinase-sensitive sequences that support cell attachment and matrix remodeling.[22] However, unmodified gelatin is thermoresponsive and lacks sufficient stability under physiological conditions, which limits its direct use for the fabrication of dimensionally stable three-dimensional (3D) scaffolds. Chemical functionalization with photocrosslinkable moieties has therefore become a key strategy to convert gelatin into a processable and cell-instructive biomaterial platform.[22–24]

### Gelatin methacryloyl

(GelMA) is currently the benchmark photocrosslinkable gelatin derivative for 3D bioprinting and soft TE.[22,25] Its frequent use originates from its straightforward synthesis, tunable degree of functionalization, compatibility with water-based processing and ability to form covalent networks upon light exposure in the presence of a photoinitiator. GelMA-based inks have been used in a wide range of soft TE contexts, including cell-laden materials, microengineered tissues and *in vitro* disease models.[25–27] Nevertheless, GelMA crosslinking proceeds mainly through radical-mediated chain-growth polymerisation, which can generate heterogeneous kinetic chains and network defects.[28,29] Norbornene-based **thiol-ene photopolymerisation** is also radical-mediated, but proceeds through a step-growth mechanism in which ring strain enhances norbornene reactivity towards thiyl radicals, supporting efficient gelation, more homogeneous network formation and improved spatiotemporal control.[28,29] In this context, norbornene-functionalized gelatin (GelNB) can be crosslinked with thiolated crosslinkers such as dithiothreitol (DTT), polyethylene glycol dithiol (PEG2SH) or thiolated gelatin (GelSH).[29,30] The choice of thiolated crosslinker provides an additional design handle, since small bifunctional crosslinkers and macromolecular gelatin-based crosslinkers can result in different crosslinking kinetics, swelling behavior, mechanical properties and biological responses.

### Digital light processing

(DLP) is particularly relevant for photocrosslinkable gelatin precursor formulations because it converts a liquid, cell-containing bioink into a covalently crosslinked hydrogel by patterned light exposure. In contrast to nozzle-based printing, the formulation is not extruded through a needle, which reduces shear-related stress during cell encapsulation. At the same time, DLP requires careful optimization of precursor concentration, photoinitiator content, light dose and absorber concentration, since these parameters determine curing depth, feature fidelity, printing time and cell compatibility.[31,32] For gelatin-based light-processing systems, feature sizes in the tens-of-micrometres range have been reported under optimized conditions. For example, a visible-light stereolithography study using GelMA reported a printing field of 9.6 × 5.4 cm² and a minimum feature size of approximately 50 µm.[33] More recently, norbornene-functionalized gelatin has been applied in DLP bioprinting of soft, cell-laden hydrogel networks. Using 2-5 wt% GelNB crosslinked with PEG4SH, Duong and Lin obtained hydrogels with G′≈120-4000 Pa, perfusable channels, > 90% viability of encapsulated human umbilical vein endothelial cells, using exposure times of 10-15 s to balance printing fidelity and cell viability.[34] These studies illustrate that gelatin-based DLP should be evaluated not only by minimum feature size, but also by the combined balance between network formation, mechanical relevance, shape fidelity and cytocompatibility.

Despite the progress in photocrosslinkable gelatin chemistry, there remains a need for modular precursor libraries that link gelatin functionalization, crosslinker selection, hydrogel network properties, DLP printability and cell response. Here, we report a small library of photocrosslinkable gelatin-based precursor formulations for cell-laden DLP, with the broader aim of generating a versatile platform for soft TE rather than a formulation restricted to one tissue model. A bifunctional methacryloyl-norbornene gelatin derivative (GelMANB) was synthesized to combine chain-growth methacryloyl crosslinking with thiol-ene norbornene chemistry and was benchmarked against GelMA. GelMANB formulations were crosslinked using different thiolated crosslinkers, including DTT and GelSH, to investigate how crosslinker chemistry as well as polymer concentration affect crosslinking kinetics, swelling, gel fraction and mechanical properties. Based on this physico-chemical screening, selected formulations were processed via DLP and evaluated for shape fidelity and cytocompatibility after encapsulation of fibroblasts. By linking gelatin chemistry, network formation, printability and cell response, this work establishes a rational workflow for selecting gelatin-based bioinks for cell-laden DLP towards soft TE platforms.

## 2. Experimental

### 2.1 Materials

Gelatin type B, isolated from bovine hides via an alkaline process, was supplied by Rousselot (Ghent, Belgium). Methacrylic anhydride (MeAnH), tartrazine (T0388), DL-dithiothreitol (DTT) (D9779) were obtained from Sigma-Aldrich (USA). Ethylenediaminetetraacetic acid tetrasodium salt tetrahydrate (EDTA·4 H_2_O), 1-(3-dimethylaminopropyl)-3-ethylcarbodiimide hydrochloride (EDC·HCl), N-hydroxysuccinimide (NHS), Millipore®Steriflip® Vacuum tube top filters (SCGP00525) were purchased from Merck (Germany). Deuterium oxide (D_2_O) was obtained from Eurisotop (Saint-Aubin Cedex, France). Spectra/Por®4 dialysis membranes (Molecular weight cut-off (MWCO) of 12,000-14,000 Da) were obtained from Spectrum Chemical Mfg. Corp. (New Brunswick, USA). SpeedCure TPO-L (ethyl (2,4,6-trimethylbenzoyl) phenyl phosphinate) was purchased from Lambson (West Yorkshire, UK). Tris(2-carboxyethyl) phosphine hydrochloride (TCEP) (T1656) was supplied by TCI Europe NV (Belgium). Fetal bovine serum (FBS) (F7524), Trypsin-EDTA solution (T4049) were supplied by Merck (USA). Calcein-AM (C1430), propidium iodide (PI) (P1304MP), and FluoroBrite™ DMEM (A1896701), ActinGreen™ 488 ReadyProbes® Reagent (R37110), 4′,6-diamidino-2-phenylindole (DAPI) (62248) were purchased from ThermoFisher Scientific (USA). Gibco™ DMEM, high glucose GlutaMAX™ (15440544), Gibco™ Antibiotic-Antimycotic (ABAM) (15240096), Dulbecco’s phosphate-buffered saline (without calcium and magnesium) (DPBS) (14190094) were obtained from Gibco (USA). MTS Assay Kit (ab197010) was obtained from Abcam (UK). Thiol-polyethylene glycol (PEG)-thiol (SH-PEG-SH) with molecular weight 3400 g·mol-1 was supplied by Laysan Bio Inc.(USA).

### 2.2 Development of photocrosslinkable gelatin derivatives

#### 2.2.1 Development of GelMA

GelMA (targeted degree of substitution (DS): 95-100%) was synthesized following the procedure described by Parmentier *et al*.[35]. First, 100 g of gelatin type B was dissolved in 1 L phosphate buffer at 40°C under stirring. After complete dissolution, the desired amount of methacrylic anhydride (2.5 eq with respect to the primary amine functionalities from ornithine and (hydroxy)lysine in gelatin type B, corresponding with 14.34 mL for 100 g; 0.3693 mmol amines/g of gelatin type B) was slowly added. After vigorously stirring for 1 hour, 1 L double distilled water was added to the reaction mixture. This mixture was then dialysed (using dialysis membranes 12-14 kDa) in reverse osmosis (RO) water for 24 hours at 40°C, while the water was changed 5 times. Lastly, the pH was adjusted to 7.4, after which it was either poured into petri-dishes or sterilized through filtration using Millipore®Steriflip® Vacuum tube top filter (pore size 0.22 µm), followed by freezing at -20°C and subsequent lyophilization, using a Christ Alpha2 freeze-dryer. The reaction schemes for the gelatin derivatives can be found in the supplementary information (Supp. Info, Figure S1).

#### 2.2.2 Development of GelSH

Thiolated gelatin (GelSH) was synthesized according to a protocol described earlier by Pien *et al.*[36], by preparing a 200 mL carbonate buffer (pH 10) and heated to 40°C. Under argon atmosphere and constant stirring, 20 g of gelatin type B was added. After formation of a homogeneous gelatin solution, EDTA (1.5 mM, tetra sodium salt tetrahydrate) was introduced. Subsequently, N-acetyl-homocysteine thiolactone was added in excess, with the amount adjusted according to the desired DS (5 eq. relative to the primary amine groups of gelatin to target a DS of 65%, 36.93 mmol). The reaction mixture was stirred vigorously at 40°C for 3 hours under inert conditions.

Following the reaction, the solution was diluted with 200 mL double-distilled water, transferred into dialysis membranes (12-14 kDa), and dialyzed against RO water at 40 °C under argon. After 24 hours of dialysis, during which the water was replaced at least five times, the pH was adjusted to 7.2-7.4. A portion of the GelSH was frozen in liquid nitrogen by slow, dropwise addition, while the remainder was reserved for sterilization and sterilized by filtration using Millipore®Steriflip® Vacuum tube top filter (pore size 0.22 µm). The material was stored at -80 °C, freeze-dried, and kept at -80 °C until further use.

#### 2.2.3 Development of GelMANB

The development of GelMANB consists of a two-step, one-pot synthesis (targeted DS MA 60-85% and DS NB 15-40%). First, 5-norbornene-2-carboxylic acid (0.70 eq with respect to the amine functionalities in gelatin type B, 12.93 mmol) was activated using EDC (0.525 eq, 9.70 mmol) and NHS (0.525 eq, 9.70 mmol) in 250 mL of DMSO. The reaction was carried out under inert argon atmosphere at room temperature for 25 hours. Simultaneously, 50 g of gelatin type B was dissolved in 750 mL dry DMSO at 40 °C. After complete dissolution, methacrylic anhydride (0.75 eq, 13.84 mmol) was added under argon, 1 hour before finishing NB activation. After vigorously stirring for 1 hour, the activated 5-norbornene-2-carboxylic acid solution was added to the gelatin mixture, which was then stirred overnight (16 h) at 50°C under argon and protected from light.

Next, the reaction mixture was precipitated in a tenfold (v/v) excess of acetone at room temperature and filtered on a glass filter (pore size 4). The resulting precipitate was then redissolved in double distilled water at 40°C and transferred in dialysis membranes (12-14 kDa). Dialysis took place for 24 hours at 40°C, while the water was changed 5 times. The pH was subsequently adjusted to 7.2-7-4, followed by pouring in petri dishes or sterilization by filtration using Millipore®Steriflip® Vacuum tube top filter (pore size 0.22 µm), freezing and subsequent lyophilization.

### 2.3 Material characterization of the modified gelatins

#### 2.3.1 ^1^H-NMR spectroscopy

To quantify the degree of substitution (DS) of the modified gelatins, proton nuclear magnetic resonance (^1^H-NMR) spectroscopy was used. Measurements were conducted using a Bruker Avance 500 MHz NMR spectrometer at 40 °C, utilizing deuterium oxide (D₂O) as the solvent. The DS is calculated by integrating the characteristic proton signals specific to the introduced functional groups - namely the methacryloyl peaks at 5.5 and 5.75 ppm or the norbornene peaks at 6.0, 6.2 and 6.3 ppm - relative to a reference peak at 1.0 ppm, corresponding to the methyl protons of the amino acids leucine (Leu), isoleucine (Ile), and valine (Val). All spectra were processed and analysed using Topspin software, applying a baseline correction to ensure accurate integration. An example of the NMR spectrum for GelMANB and the indicated peaks can be found in supplementary information (Supp. Info, Figure S2). The DS was calculated based on the following formulas:

GelMA:

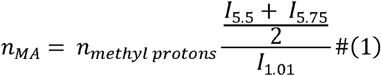

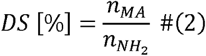

With:

I_d= 5.5–5.75_ ppm = □integral of the signal characteristic for the protons of methacrylamide

I_d= 1.0_ ppm = □integral of the signal of the protons of the reference peak

*n_NH2_*= □0.03693 moles primary amines per 100 g gelatin type B

*n_methyl protons_*=□ 0.384 moles methyl protons (in reference signal) of Val, Leu and Ile per 100 g gelatin type B

GelNB:

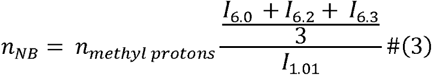

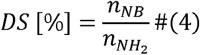

With:

I_d= 6.3-6.0_ ppm = □integral of the signal characteristic for the protons of norbornene I_d= 1.0_ ppm = integral of the signal of the protons of the reference peak

*n_NH2_*=□ 0.03693 moles primary amines per 100 g gelatin type B

*n_methyl protons_*=□ 0.384 moles methyl protons (in reference signal) of Val, Leu and Ile per 100 g gelatin type B

#### 2.3.2 Ortho-phthalic dialdehyde (OPA) assay

The OPA assay was used to determine the DS of GelSH.[36] In brief, a GelSH sample was reacted with OPA and 2-mercaptoethanol, after which the absorbance was recorded at 335 nm using UV-VIS spectroscopy, with a blank sample as reference. A calibration curve generated from n-butylamine standards with known amine concentration (0.002 M-0.01 M) was used to calculate the DS based on the amount of unreacted amine groups. All measurements were performed in triplicate.

#### 2.3.3 Synthesis of photo-initiator and resin preparation

Lithium phenyl-2,4,6-trimethylbenzoyl phosphinate (LAP) was synthesized as described in [37].

To prepare the resins, the photocrosslinkable gelatin precursors were dissolved in double-distilled water at 40°C to obtain solutions with three different gelatin concentrations: 5, 7.5 and 10 w/v%. For formulations containing both GelMANB and GelSH, these concentrations refer to the total gelatin concentration, corresponding to the combined mass of GelMANB and GelSH. In contrast, for GelMANB/DTT and GelMANB/PEGSH formulations, the stated gelatin concentration refers only to the GelMANB content, since DTT and PEGSH do not contribute to the gelatin mass fraction. After dissolution, LAP was added to a final concentration of 2 mol% (relative to the number of photocrosslinkable functionalities in the corresponding gelatin precursors). For formulations containing thiolated crosslinkers, TCEP was added at 25 mol% relative to thiol functionalities to limit disulfide formation before photocrosslinking. The same TCEP concentration was also included in GelMANB formulations without thiolated crosslinker as a reference condition, allowing comparison with the thiol-containing GelMANB resins under comparable additive conditions. DTT was added from a freshly prepared 80 mg·mL^-1^ stock solution, whereas PEGSH and GelSH were added directly to the resin mixture after dissolution of all other components. DTT, PEGSH and GelSH were added in an equimolar ratio of thiol-to-ene functionalities. These mixtures were then used for further characterization.

#### 2.3.4 *In situ* crosslinking using photorheology

To assess the crosslinking dynamics of the different resins, photorheology was performed with an Anton Paar Physica MCR 301 rheometer, equipped with an Omnicure S1500 UV-A light source, featuring a filter for wavelengths between 400-500 nm. For each measurement, 270 µL of gelatin precursor solution containing 2 mol% LAP was introduced between two parallel plates using a 25 mm spindle and a 0.3 mm gap. The storage modulus G′ and loss modulus G″ were recorded at 37 °C, 0.1% strain and 1 Hz. The measurement consisted of three consecutive intervals: 1 min before irradiation, 9 min during UV-A irradiation at 20.1 mW·cm^-2^, and 5 min after irradiation, resulting in a total measurement time of 15 min. All measurements were performed in triplicate (n = 3).

#### 2.3.5 Formation of crosslinked networks through film casting

Resin formulations were prepared as described in 2.3.3. The heated solution was poured between two parallel glass plates covered with release foil (Holders Technology, ACC-3) and separated by a 1 mm thick silicone spacer. Then, the material was placed in the fridge for 30 min at 4°C for physical gelation and subsequently irradiated from both sides (top and bottom) with UV-A light (λ = 365 nm, 8 mW·cm^−2^) for 30 min.

#### 2.3.6 Evaluation of the gel fraction and mass swelling ratio

Cylindrical samples (diameter: 7 mm, thickness: 1 mm, n=5) were punched from the film cast sheets of GelMA and GelMANB (preparation described in section 2.3.5). For gel fraction assessment, the samples were freeze-dried and weighed in dry state to determine the dry mass (*m_d1_*). Next, the samples were incubated in double distilled water at 37°C for 72 hours, and freeze-dried again to determine the second dry mass (*m_d2_*). The gel fraction was then determined using the following equation:

For the mass swelling ratio, the samples were allowed to equilibrium swell (37°C, 72 h) after which the mass of the swollen samples (m_s_) was quantified and subsequently freeze-dried. Then, the dry mass was determined (m_d_). The mass swelling ratio (MSR) was then determined using the following equation:

#### 2.3.7 Frequency sweep of crosslinked hydrogel samples

The viscoelastic properties of the materials were determined by a frequency sweep analysis using a Physica MCR 301 rheometer from Anton Paar. After incubating the crosslinked sheets obtained through film casting (section 2.3.3) in double distilled water at 37°C for 72 hours to reach swelling equilibrium, cylindrical samples (14 mm diameter, 1 mm thickness) were punched out. The samples were positioned between two parallel plates (15 mm spindle) and a frequency sweep analysis was performed from 0.1 to 10□Hz (F_N_ = 0.5□N, amplitude of 0.1□% and T□=□37°C) to determine the storage (G′) and loss (G′’) moduli. These measurements were performed in triplicate (n=3). The compressive modulus (E) was determined using the storage modulus via the following equation, where ν is the Poisson number, which is assumed to be 0.5 for ideal hydrogels.[38]

#### 2.3.8 Evaluation of mechanical properties through tensile testing

Rings of 14□mm inner diameter and 18□mm outer diameter with a thickness ≈□1.5□mm were punched out from the film cast sheets (after equilibrium swelling, 72h at 37°C). The tensile properties of the GelMA and GelMANB samples with different thiolated crosslinkers were determined at room temperature using a universal testing machine (Tinius Olsen 3ST) equipped with a 500□N load cell. The samples were positioned as shown in Figure 1, between two 3D-printed curved hooks with a diameter of 5□mm. This method was specifically developed for gelatin-based hydrogels by Pien *et al*.[36], as traditional dogbone-shaped hydrogel samples encounter fracture issues when clamped between the grips. Samples were subjected to mechanical testing with an initial preload force of 0.01 N at a crosshead velocity of 5 mm·min⁻ ¹. The resulting displacement and force data were used to generate stress-strain plots. From the measured displacement (Δs), the internal circumference (C) was determined using a formula incorporating the diameter of the 3D-printed hooks (d_pin_ = 5□mm):

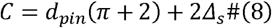

**Figure 1.**
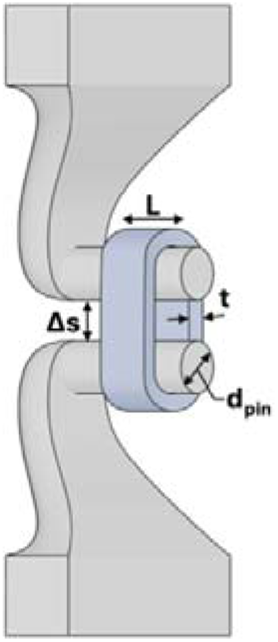
Schematic set-up for ring-shaped tensile testing.

Circumferential strain (ε) was defined as the ratio of C to the initial internal circumference (C_init_), while circumferential stress (σ) was derived from the applied load (F) normalized by the specimen’s geometry - specifically its length (L) and wall thickness (t). The Young’s modulus was then extracted from the slope of the initial linear region of each stress-strain curve.

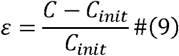

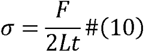

### 2.4 Material processing using digital light processing (DLP)

#### 2.4.1 Polymer and printing parameter optimization

For the optimised formulations, GelMA (10 w/v%) and GelMANB (10 w/v% and 7.5 w/v%) were dissolved in Dulbecco’s phosphate-buffered saline (DPBS, pH 7.0–7.3) at 40 °C. After complete dissolution, LAP and tartrazine stock solutions were added to obtain final concentrations of 10 mol% LAP and formulation-specific tartrazine concentrations of 2.0 mol% for GelMA-based resins and 2.44 mol% for GelMANB-based resins, relative to the number of photocrosslinkable functionalities present. The LAP concentration was selected based on the study by Maes et al., in which 10 mol% LAP was optimised for DLP printing of GelMANB- and GelMA-AEMA-based resins.[39]

Tartrazine was selected as photoabsorber to reduce off-focal plane polymerisation and improve printing fidelity. This hydrophilic yellow dye absorbs around 405 nm and has previously been used as a biocompatible photoabsorber for light-based bioprinting.[40] Since the optimal photoabsorber concentration depends on the specific resin formulation, reported tartrazine concentrations vary substantially. For example, Duong et al. screened tartrazine concentrations between 0 and 1.5 mM, corresponding to 0-0.80 mg·mL⁻¹, for DLP printing of gelatin-norbornene hydrogels and identified 1.2 mM, corresponding to 0.64 mg·mL⁻¹, as optimal.[41] Based on this range, tartrazine concentrations between 0.5 and 3 mol% relative to the number of photocrosslinkable functionalities were screened for the GelMA- and GelMANB-based resins in the present study. Concentrations above 2.5 mol% prevented efficient gel formation, whereas concentrations below 2 mol% resulted in overcuring and partial or complete loss of the porous scaffold architecture. Based on this formulation-specific screening, 2.0 mol% tartrazine was selected for GelMA-based resins, while 2.44 mol% tartrazine, corresponding to 0.5 mg·mL⁻¹, was selected for GelMANB-based resins.

For formulations containing thiolated crosslinkers, neutralised TCEP was added at 25 mol% relative to the thiol functionalities to limit premature disulfide formation. The thiolated crosslinker, either DTT or GelSH, was subsequently added as the final component using an equimolar thiol-to-ene ratio. After complete dissolution and sonication for 5 min, the resin was transferred to the vat of the DLP printer (CELLINK Lumen-X, CELLINK, BICO company, USA). The final resin formulations and printing conditions selected based on photoabsorber screening, working curve analysis and scaffold printability are summarised in Table 1.

**Table 1.**
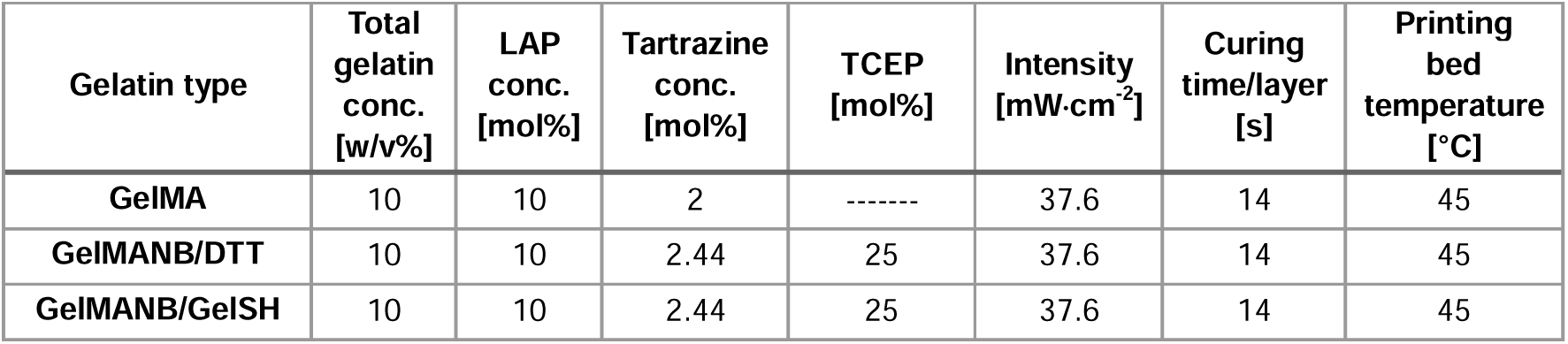
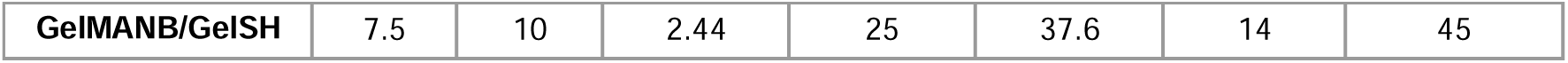
Overview of the resin formulations and printing parameters.

Working curves were established for the selected resin formulations by printing 1 cm² square samples at different exposure times using a nominal layer thickness of 100 µm. The light intensity was set to 37.6 mW·cm⁻² and the resolution to 100 µm for all resins. The cured layer thickness was measured using a micrometer after subtraction of the glass coverslip thickness. Three samples were measured per exposure time for each material (n = 3). The working curves are presented in Figure S3. The penetration depth (D_p_) and critical energy dose required for polymerisation (E_c_) were determined by fitting the resulting data to the working curve equation:

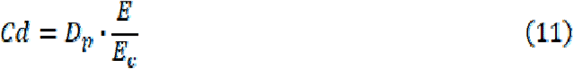

where C□ is the cured layer thickness, D□ is the penetration depth of the curing light, E is the applied light dose and E is the critical energy dose required for polymerisation. The minimal exposure time required for polymerisation (t_c_) was subsequently calculated from E_c_ and the applied light intensity.

Various structures were printed for further evaluation. An overview of the CAD images, including their dimensions, is shown in Figure 2. CAD models were designed using Tinkercad (Autodesk Inc., USA).

**Figure 2.**
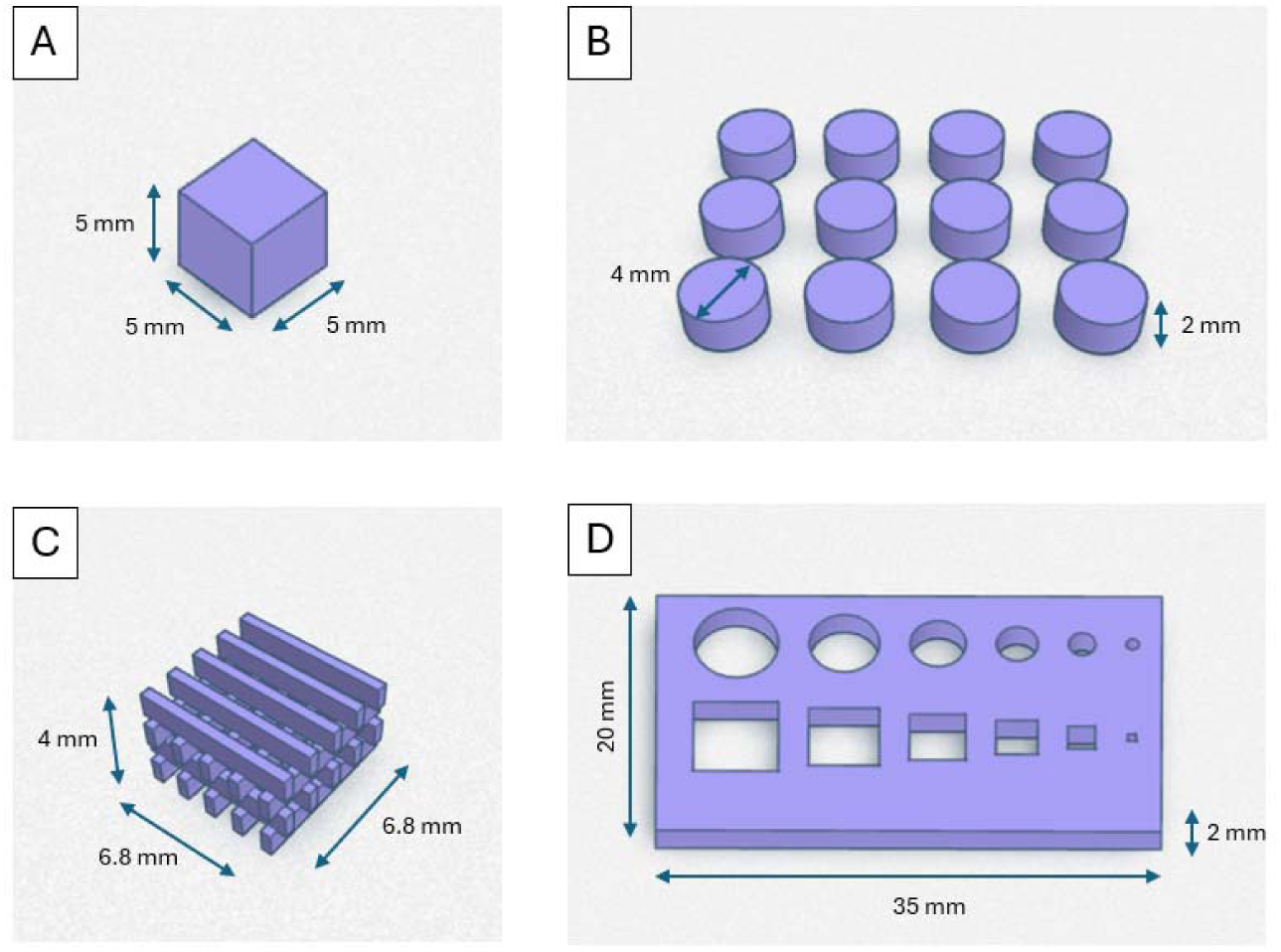
CAD designs. (A) solid cube for gel fraction and swelling ratio assessment (5 mm x 5 mm x 5mm); (B) discs for MTS assay (8 discs with 4mm diameter and 2 mm height); (C) TE scaffolds consisting of a cube with struts and pores (0.4 and 1.3 mm, respectively); (D) benchmark design to assess the resolution of the optimized parameters (6 different sizes for length of square and diameter of circle going from 6 to 1 mm).

#### 2.4.2 Morphological and physico-chemical evaluation of DLP printed constructs

A porous, cubic 3D scaffold and a benchmark were printed to obtain information regarding the computer-aided design/computer-aided manufacturing (CAD/CAM) mimicry using a light microscope (Figure 2 C,D). After printing, scaffolds were washed in warm ultrapure water (UPW) and immersed in 10x PBS to densify the network for imaging. Increasing the ionic strength of the surrounding medium is known to generate an outward osmotic water gradient that lowers the equilibrium water content and mesh size of gelatin-based hydrogels).[42] The resulting reduction in water content increased the density and opacity of the scaffolds, improving contrast for photographic imaging of their architecture. For the swelling degree and gel fraction determination, the cubic samples (CAD - Figure 2A) were frozen directly after printing. DLP-printed disc-shaped samples (Figure 2B) also underwent gel fraction (GF) and mass swelling ratio (MSR) analysis following the protocol described by Parmentier *et al.* [43], whereby samples were swollen for 24 hours.

### 2.5 DLP bioprinting and *in vitro* evaluation

#### 2.5.1 Cell lines and culture conditions

Human foreskin fibroblasts (HFF-1, ATCC, SCRC-1041) were used as a standardized stromal fibroblast model to evaluate cell viability, morphology and proliferative response following encapsulation and DLP bioprinting. Fibroblasts were selected because they are broadly relevant stromal cells involved in extracellular matrix deposition and remodeling, and fibroblast-like mammalian cells are commonly used for *in vitro* cytotoxicity screening according to ISO 10993-5. HFF-1 were cultured in DMEM high glucose supplemented with 10% of FBS, 1% ABAM (100 U/mL penicillin, 100 µg/mL streptomycin, 0.25 µg/mL of Gibco Amphotericin B). Cells were seeded in flasks at a seeding density of 0.008×10^6^ cells·cm^-2^ and cultured at 37°C in a humidified atmosphere containing 5% CO_2_. Medium was renewed 2 times per week. The cells of passages 5-8 were used in subsequent experiments.

#### 2.5.2 Fabrication of cell-laden scaffolds via DLP biofabrication

For the fabrication of cell-laden scaffolds, the cells were detached through enzymatic digestion using trypsin/EDTA. Then the cells were pelleted by centrifugation at 300 g for 10 min, at room temperature (RT). The obtained pellets were resuspended in fresh culture medium. For the introduction of the cells into the resins, the cells were again pelleted by centrifugation at 300 g for 10 min, RT. Immediately before printing, the supernatant was removed, and the cell pellet was gently resuspended in the resin to a final cell concentration of 2·10^6^ cells·mL^-1^. The cell-containing resins were then used for DLP printing. After printing, the scaffolds were washed with warm DPBS, transferred into the corresponding culture medium, and cultured for 2-7 days at 37°C, in a humidified atmosphere containing 5% CO_2_. The culture medium was exchanged 18-24 hours post-printing, and then every 2-3 days.

#### 2.5.3 Metabolic activity of cells in DLP-printed scaffolds

The metabolic activity of the cells, encapsulated into DLP-printed scaffolds, was determined using MTS assay on day 2, 5, and 7 post-printing. For the assay, the disk scaffolds were transferred to the wells of a 96-well plate and 120 µL of fresh culture medium containing 20 µL of MTS reagent was added per well. Culture medium containing 20 µL of MTS reagent (in wells without cells) served as the negative control. The scaffolds were incubated at 37°C in a humidified atmosphere containing 5% CO_2_ for 2 hours. To avoid the interference of the scaffolds with spectrophotometric detection of the MTS-conversion product, we transferred the resulting culture medium into new wells of a 96-well plate. The absorbance was measured at 490 nm using TECAN Infinite M200 Pro multimode plate reader (Tecan Group Ltd., USA). All conditions were performed in triplicate (n=3), and the results are presented as the mean± standard deviation (SD).

#### 2.5.4 Analysis of cell viability

To assess cell viability on day 3, 5, and 7 post-printing, DLP-printed cell-laden scaffolds were gently washed 3 times with warm DPBS to remove the residual culture medium and incubated with calcein-AM/PI solution (2 μg mL−1 of each fluorescent probe in FluoroBrite™ DMEM) for 15 min, protected from light. Images were captured using a confocal laser scanning microscope (LSM 710, Zeiss, Germany) equipped with a green fluorescent protein and a Texas Red filter. To capture cells’ distribution throughout the 3D structure of the scaffolds, Z-stack image series were acquired across the sample depth using a step size of 20 µm over a total Z-range of 500-1200 µm. The number of living/dead cells was counted on the individual optical sections (n= 14) using ImageJ software. The biocompatibility of the hydrogel discs was assessed based on morphology. When the circularity was > 0.5, the cell was classified as spherical; circularity <0.5 was classified as elongated. Elongated morphology indicates high biocompatibility and spherical morphology lower biocompatibility. Maximum intensity projections (MIP) were generated from the z-stacks using ZEN software.

#### 2.5.6. Cell morphology analysis

To assess the morphology of the cells in cell-laden scaffolds, on day 7 post-printing, the cells were fixed with 4% w/v PFA for 24 hours at 4°C, and permeabilized with 0.1% Triton-X-100 in DPBS for 20 min at RT. Upon permeabilization, the cells were washed twice with DPBS. The F-actin was stained with ActinGreen™ 488 ReadyProbes® Reagent, while the nuclei were counterstained with 4′,6-diamidino-2-phenylindole (DAPI). Cells were imaged using confocal laser scanning microscopy (LSM 710, Zeiss, Germany). Z-stack image series were acquired across the sample depth using a step size of 20 µm over a total Z-range of 847-1894 µm. Images were taken at λex = 405 nm (DAPI), λex = 488 nm (ActinGreen), and λ_em_=575 nm.

Images were analyzed in Fiji/ImageJ 2.18.0/1.54p using MorphoLibJ plugin [44]. The DAPI and F-actin channels were separated, background-corrected, and independently thresholded. DAPI-positive nuclei were separated by watershed segmentation and converted into uniquely labelled markers using connected-component labelling. An F-actin-positive cellular mask was generated by thresholding the actin channel, and background particles were excluded using Analyze Particles. Marker-controlled watershed segmentation was then performed using the labelled DAPI nuclei as markers, an actin-derived gradient image as the input image, and the binary actin mask as the restriction mask. Morphological measurements were obtained from the resulting labelled cell regions, whereas actin intensity was measured from the original actin channel. Morphology of living cells was quantified based on circularity, as described above.

### 2.6 Statistical analysis

Statistical analysis was performed using GraphPad Prism (version 11.0.2; Dotmatics, USA). Data distribution and model assumptions were assessed before statistical testing. Normality was evaluated by visual inspection of Q-Q plots, and equality of variances was assessed by plotting the residuals against the fitted values. When the analysed datasets met the assumptions for parametric testing, one-way or two-way ANOVA was used, followed by the appropriate post hoc multiple comparison test, as indicated in the corresponding figure captions. In case the analysed datasets did not meet the assumption of homogeneity of variances, group means were compared using *Brown-Forsythe and Welch’s ANOVA, followed by Dunnett’s T3 multiple comparisons test.* The statistical tests and number of replicates are indicated in the figure and table captions. Values are expressed as mean ± SD. Differences with p < 0.05 were considered statistically significant.

## 3. Results and discussion

### 3.1 Development and characterization of photo-crosslinkable gelatins

#### 3.1.1 Determination of degree of substitution

GelMANB was synthesized as a bifunctional, photocrosslinkable gelatin derivative bearing both methacrylamide and norbornene functionalities on the gelatin backbone (Supp. Info, Figure S1). This design enables the combination of chain-growth methacrylamide crosslinking with step-growth thiol-ene crosslinking, thereby providing a strategy to tune the resulting hydrogel network architecture. In this work, GelMANB with a methacrylamide degree of substitution (DS) of 66-68% and a norbornene DS of 24-28% was used throughout the experiments. A highly substituted GelMA derivative with a DS of 96-98% was included as reference material, since its total amount of crosslinkable functionalities approximated that of the GelMANB formulations. GelSH with a DS of 69% was used as macromolecular thiolated crosslinker in the GelMANB/GelSH thiol-ene system.

Successful functionalization of gelatin type B was confirmed by ¹H-NMR spectroscopy. In the GelMANB spectrum (Supp. Info, Figure S2), the characteristic methacryloyl proton signals were observed at approximately 5.5 and 5.7 ppm, while the norbornene proton signals appeared at approximately 6.0, 6.2 and 6.3 ppm. The DS was calculated by integration of these signals relative to the reference peak at 1.0 ppm, originating from the methyl protons of leucine, isoleucine, and valine residues in gelatin. This analysis confirmed that 96% of the available amine groups in GelMA was functionalized with methacrylamide groups, corresponding to 0.354 mmol MA functionalities·g⁻¹ gelatin. For GelMANB, 68% of the amine groups was functionalized with methacrylamide groups, corresponding to 0.244 mmol MA functionalities·g⁻¹ gelatin, while 28% were functionalized with norbornene groups, corresponding to 0.103 mmol NB functionalities·g⁻¹ gelatin. These results confirm the successful synthesis of a bifunctional GelMANB precursor with a total crosslinkable functionality content comparable to highly substituted GelMA, while introducing an additional norbornene handle for thiol-ene network formation.

#### 3.1.2 Evaluation of storage modulus and crosslinking kinetics via photo-rheology

The *in situ* photocrosslinking behaviour of the different gelatin precursor formulations was evaluated by photorheology to assess both crosslinking kinetics and final network stiffness. The initial increase in storage modulus, G′, provides an indication of the crosslinking rate, which is relevant for cell-laden light-based processing because rapid network formation helps stabilise encapsulated cells within the printed geometry while limiting the required light exposure. In contrast, the plateau value of G′ reflects the final stiffness of the formed hydrogel network and therefore provides insight into the mechanical range accessible for soft tissue engineering applications. The influence of total gelatin concentration (w/v%) and thiolated crosslinker type was therefore investigated for GelMA and GelMANB-based formulations (Figure 3).

**Figure 3.**
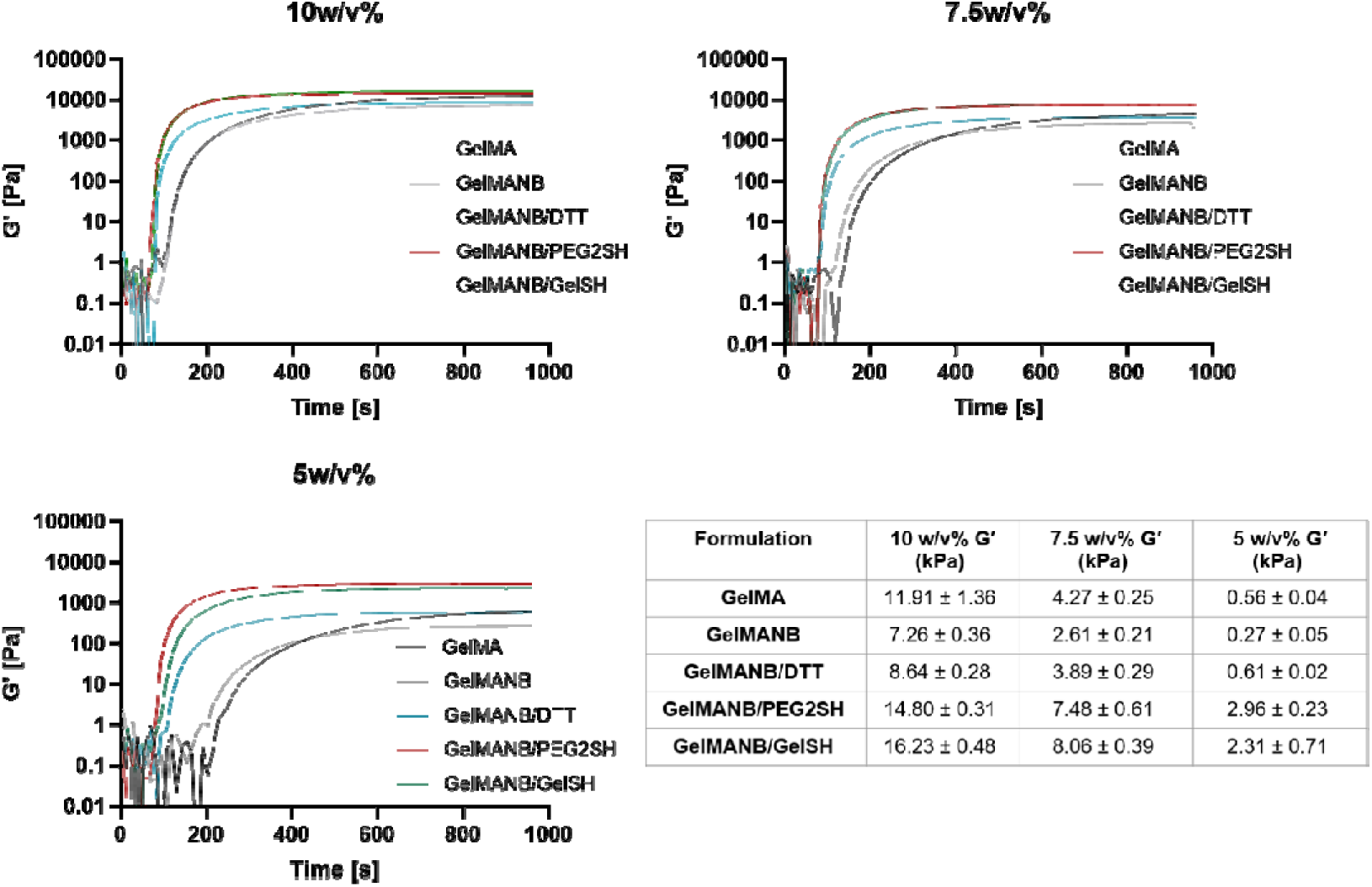
Photorheological evaluation of GelMA and GelMANB-based precursor formulations. Storage modulus (G′) as a function of time for formulations prepared at 10 w/v%, 7.5 w/v% and 5 w/v% total gelatin concentration, with or without thiolated crosslinkers, including DTT, PEG2SH and GelSH. The table summarizes the mean plateau G′ values for the 10 w/v% and 7.5 w/v% formulations.

The formulations covered a broad range of storage moduli, from 0.23±0.01 kPa for 5 w/v% GelMANB without additional thiolated crosslinker to 16.23±0.02 kPa for 10 w/v% GelMANB/GelSH. As expected, increasing the total gelatin concentration (5 w/v% to 10 w/v%) resulted in higher G′ values, confirming that polymer content is a key parameter for tuning hydrogel stiffness. At 10 w/v%, the highest G′ values were obtained for GelMANB/GelSH and GelMANB/PEG2SH, reaching 16.23±0.02 kPa and 14.80±0.01 kPa, respectively, followed by GelMA (11.85±0.38 kPa), GelMANB/DTT (8.63±0.02 kPa) and GelMANB without thiolated crosslinker (7.24±0.14 kPa). At 7.5 w/v%, the same crosslinker-dependent trend was observed, with GelMANB/PEG2SH and GelMANB/GelSH reaching 8.90±0.02 kPa and 8.06±0.02 kPa, respectively, while GelMA, GelMANB/DTT and GelMANB reached 4.24±0.23 kPa, 3.88±0.02 kPa and 2.61±0.17 kPa, respectively.

Several formulations therefore reached storage moduli within the low-kPa to tens-of-kPa range relevant for soft tissue engineering, while still allowing straightforward handling. In particular, 10 w/v% GelMANB/DTT, 7.5 w/v% GelMANB/PEG2SH and 7.5 w/v% GelMANB/GelSH resulted in G′ values within or close to the range targeted for compliant soft tissue models (e.g. alveolar tissue (∼4-8 kPa)). At 10 w/v%, GelMA displayed a significantly higher G′ compared to GelMANB/DTT (p < 0.0001). This difference may be related to the higher methacryloyl content of GelMA and the resulting chain-growth contribution to network formation, whereas GelMANB/DTT relies more strongly on norbornene-mediated thiol-ene crosslinking through a small bifunctional crosslinker.[45] In contrast, the final stiffness of the GelMANB-based networks was strongly dependent on the selected thiolated crosslinker. PEG2SH and GelSH resulted in comparable plateau moduli, indicating that both polymeric and macromolecular thiolated crosslinkers contributed effectively to network formation. For PEG2SH, this may be related to its flexible polymeric spacer, which can bridge norbornene-functionalized gelatin chains over a longer distance. For GelSH, the macromolecular and multifunctional gelatin backbone provides multiple thiol groups and additional gelatin content, which can increase the number of elastically effective network connections. DTT consistently resulted in lower G′ values, which is in line with previous GelNB thiol-ene studies from our group showing that the architecture of the thiolated crosslinker, including molecular weight, functionality and macromolecular character, strongly affects hydrogel network properties and printability.[29] As a small bifunctional crosslinker, DTT can form covalent connections between norbornene-functionalized gelatin chains, but lacks the longer spacer length and multifunctional macromolecular character of PEG2SH and GelSH, which may reduce its contribution to elastically effective network formation.

The crosslinking rate, calculated from the slope of the linear region of the G′ increase, further confirmed the strong influence of crosslinker chemistry (Figure 4). GelMANB-based thiol-ene formulations displayed faster crosslinking compared to GelMA, consistent with the rapid step-growth thiol-ene reaction. This is attributed to ring-strain relief following thiyl radical addition to the norbornene group and the rapid subsequent hydrogen abstraction from a thiol, which regenerates the thiyl radical and propagates the reaction.[29] Among the GelMANB formulations, GelMANB/GelSH and GelMANB/PEG2SH showed markedly higher slopes than GelMANB/DTT, indicating faster network formation. This faster increase in G′ is relevant for digital light processing, where rapid gelation can reduce the time during which scattered light may induce crosslinking outside the intended region.[46] Overall, the photorheology results demonstrate that the GelMANB platform enables crosslinker-dependent tuning of both hydrogel stiffness and crosslinking kinetics, with GelSH and PEG2SH providing the most rapid network formation and highest final storage moduli.

**Figure 4.**
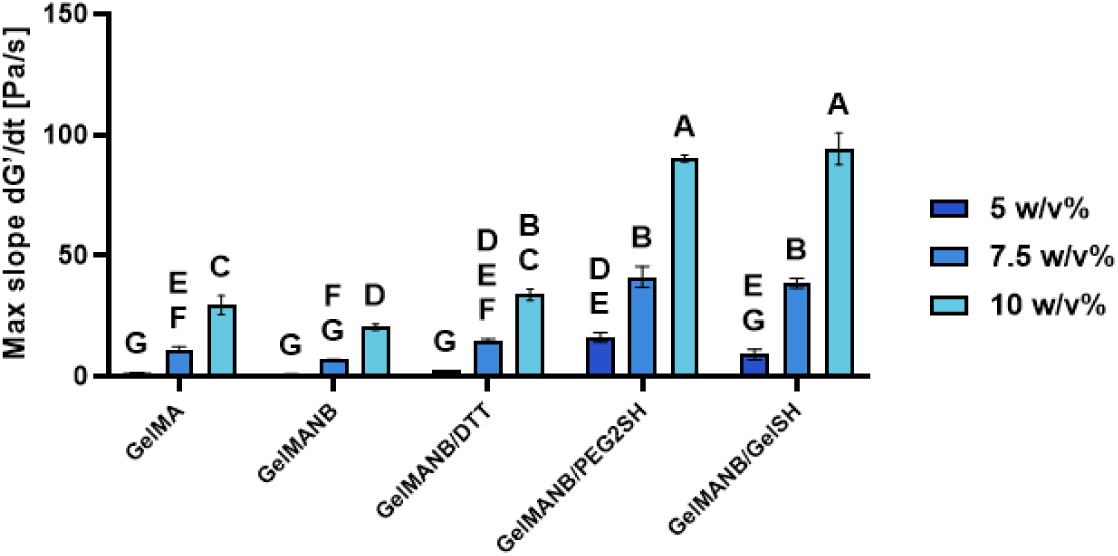
Crosslinking rate of 10, 7.5 and 5 w/v% GelMA and GelMANB-based formulations, calculated from the slope of the linear region of the photorheological G′ increase. Statistical analysis was performed using two-way ANOVA with Bonferroni’s multiple comparison test. Different letters indicate significant differences at p < 0.05. Statistical significance indicates differences in crosslinking kinetics between formulations.

#### 3.1.3 Viscoelastic behaviour of equilibrium-swollen hydrogel networks

The viscoelastic behaviour of the crosslinked hydrogel networks was evaluated by frequency sweep analysis on equilibrium-swollen 10 w/v% hydrogel films at 37 °C. Since cell-laden hydrogels are ultimately cultured in a hydrated environment, these measurements provide insight into the mechanical behaviour of the networks under more application-relevant conditions. The storage modulus (G′) and loss modulus (G″) were recorded over a frequency range of 0.1-10 Hz under a constant normal force of 0.5 N.

For all formulations, G′ remained relatively constant across the investigated frequency range, indicating the formation of stable crosslinked hydrogel networks. Moreover, G′ was consistently higher than G″ (Figure 5A), and no crossover between the two moduli was observed over the entire frequency range. This confirms that all formulations exhibited predominantly elastic, solid-like behaviour under the applied oscillatory conditions. The weak frequency dependence of both G′ and G″ further suggests that the crosslinked networks maintained their structural integrity and did not undergo significant relaxation or rearrangement within the tested frequency range.

**Figure 5.**
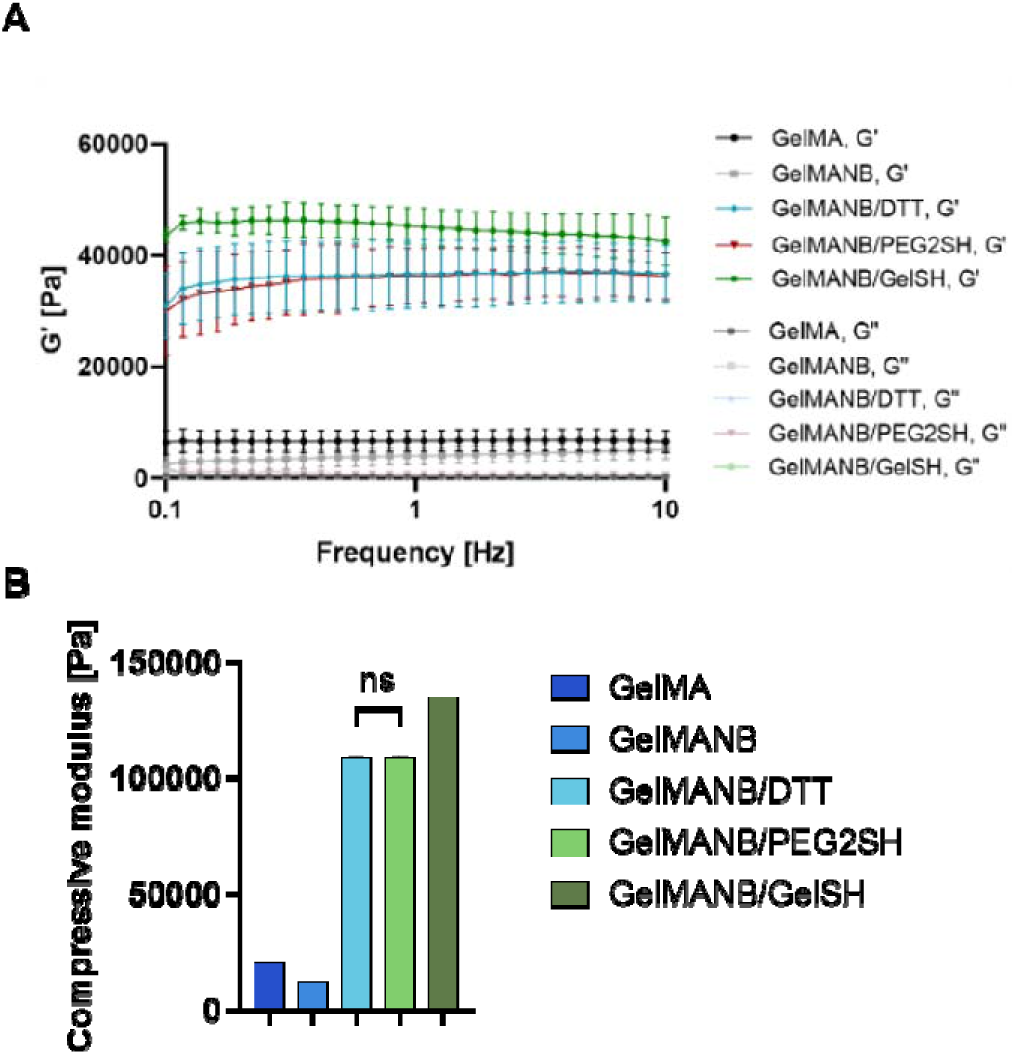
(A) Frequency sweep analysis of equilibrium-swollen, film-cast hydrogels prepared at 10 w/v% total gelatin concentration. Storage modulus (G′) and loss modulus (G″) are shown as a function of frequency from 0.1 to 10 Hz. (B) Compressive storage modulus calculated from frequency sweep data using E′ = 2G′(1 + ν), assuming a Poisson’s ratio of 0.5. Statistical analysis was performed using one-way ANOVA with Tukey’s post hoc test. Significant differences were observed between all groups (p < 0.0001), except for GelMANB/DTT versus GelMANB/PEG2SH

The magnitude of G′ was strongly dependent on the crosslinking chemistry. GelMANB formulations containing thiolated crosslinkers displayed substantially higher G′ values than GelMA or GelMANB without additional thiolated crosslinker. In particular, GelMANB/GelSH showed the highest storage modulus, followed by GelMANB/PEG2SH and GelMANB/DTT, whereas GelMANB without thiolated crosslinker resulted in the lowest values. This trend confirms that the incorporation of thiolated crosslinkers into GelMANB networks substantially reinforces the hydrogel structure, with the macromolecular GelSH crosslinker providing the most pronounced effect.

Using G′ values obtained from the frequency sweep analysis, the compressive storage modulus (E′) was calculated according to E′ = 2G′(1 + ν), assuming a Poisson’s ratio of 0.5 for ideal hydrogels.[38] The resulting compressive moduli are displayed in Figure 5B. GelMANB without thiolated crosslinker resulted in the lowest compressive modulus (11.96±0.13 kPa), consistent with the lower number of effective crosslinks in the network. In contrast, GelMANB/GelSH reached the highest compressive modulus (134.75±0.22 kPa), while GelMANB/DTT and GelMANB/PEG2SH showed comparable values, with no significant difference between these two groups. Significant differences were observed between all other groups (p < 0.0001).

These values are in line with, and extend, the mechanical range reported for related gelatin-based photocrosslinked hydrogels. In the VAM study by Pien and Bogaert *et al.*, GelNB/GelSH formulations showed storage moduli ranging from approximately 0.2 to 12.5 kPa, depending on formulation and crosslink density.[36] Van Damme *et al.* [12] reported G′ values of 14.0±2.8 kPa for chemically crosslinked GelMA100, 10.1±1.4 kPa for GelMA65, 9.3±1.9 kPa for GelNB55/SH75 and 17.9±2.1 kPa for GelNB89/SH75. After combined physical and chemical crosslinking, G′ increased to 53.3-64.1 kPa for GelMA and 24.1-31.1 kPa for GelNB/SH films.[12] Parmentier *et al.* further reported compressive moduli E’ of 76.09±1.68 kPa for 10 w/v% GelMA 2.5 eq., 85.60±11.96 kPa for 10 w/v% GelNB 2.5 eq. crosslinked with GelSH, and 111.05±7.85 kPa for 10 w/v% GelNBNB/GelSH networks.[43] In a later GelMA structure-function study, Parmentier *et al.* reported lower E′ values for GelMA-only networks, with 1.73 kPa for 10 w/v% GelMA34, 4.42 kPa for 5 w/v% GelMA96, and 10.18 kPa for 10 w/v% GelMA96.[35] The E′ range obtained in the present study, from 11.96±0.13 kPa to 134.75±0.22 kPa, therefore overlaps with and slightly extends the upper range previously reported for gelatin-based methacryloyl and thiol-norbornene networks. In particular, the high E′ of GelMANB/GelSH is consistent with the reinforcing effect of combining norbornene-mediated thiol-ene crosslinking with a macromolecular, multifunctional GelSH crosslinker. Overall, the frequency sweep results show that thiolated GelMANB networks exhibit enhanced elastic and compressive properties compared to GelMA and non-thiolated GelMANB, supporting the role of thiol-ene crosslinking and crosslinker architecture in reinforcing the hydrogel network.

#### 3.1.4 Determination of swelling properties and gel fraction

Based on the photorheology and frequency sweep results, the 10 w/v% formulations were selected for further physicochemical characterisation because they provided stable hydrogel networks with clearly distinguishable mechanical properties for each crosslinking chemistry, while still allowing straightforward handling and comparison between GelMA and GelMANB-based systems. The gel fraction and mass swelling ratio of the 10 w/v% hydrogel films were determined to evaluate network formation and hydration behaviour (Figure 6). The gel fraction provides an indication of the amount of insoluble, covalently crosslinked material remaining after extraction and therefore reflects the efficiency of network formation. This is particularly relevant for cell-laden applications, since soluble or unreacted precursor components may leach from the hydrogel and affect cytocompatibility.

**Figure 6.**
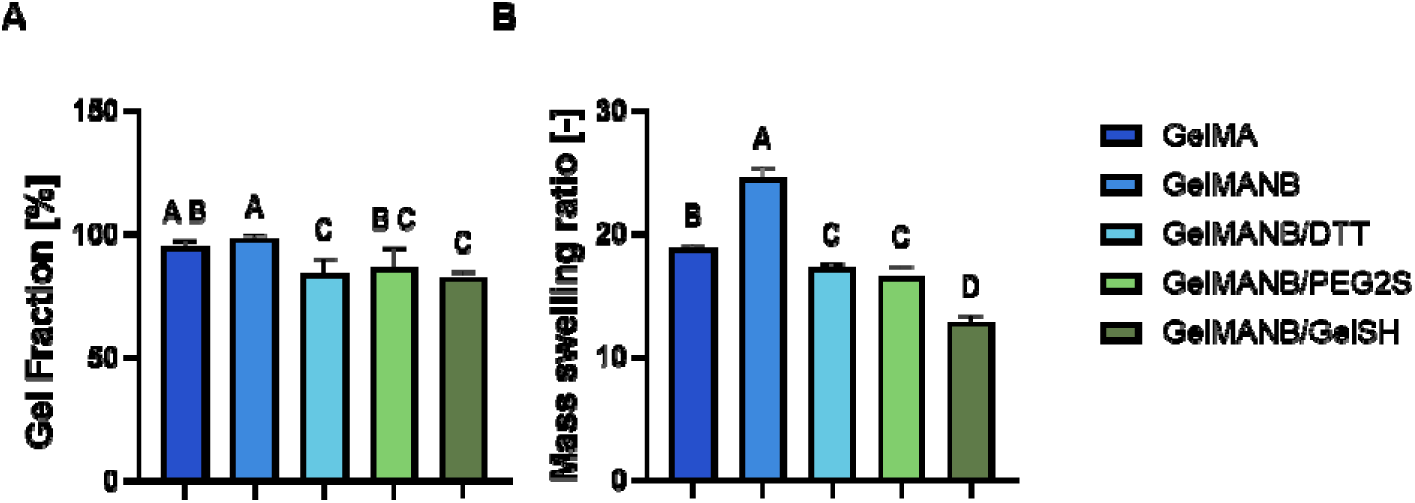
Gel fraction (A) and mass swelling ratio (B) of 10 w/v% GelMA and GelMANB-based hydrogel films. Gel fractions were determined after extraction of soluble components, while mass swelling ratios were calculated from the swollen and dry masses of the hydrogels. Statistical analysis was performed using one-way ANOVA with Tukey’s post hoc test. Different letters indicate significant differences at p < 0.05.

All hydrogel formulations tested showed high gel fractions, exceeding 84%, indicating the formation of stable hydrogel networks with limited leachable content, in line with previous reports on GelMA, GelNB/DTT and GelNB/GelSH systems.[47] The mass swelling ratio was used to assess the water uptake capacity of the crosslinked networks. This parameter is influenced by polymer volume fraction, polymer-solvent interactions, molar mass between crosslinks and overall crosslinking density.[48] It also gives an indication of the ability of the hydrogels to mimic the aqueous cellular environment of the ECM.[49]

GelMANB without additional thiolated crosslinker showed the highest mass swelling ratio (24.5±0.95), followed by GelMA (18.8±0.3). Incorporation of thiolated crosslinkers reduced the swelling ratio to 17.26±0.34 for GelMANB/DTT, 16.52±0.84 for GelMANB/PEG2SH and 12.87±0.49 for GelMANB/GelSH. This decrease in swelling upon addition of thiolated crosslinkers indicates that the hydration behaviour of GelMANB-based networks is strongly governed by thiol-ene network formation and crosslinker architecture. The lower swelling of GelMANB/DTT and GelMANB/PEG2SH compared to GelMANB without additional crosslinker suggests more efficient network formation through covalent thiol-ene crosslinking, while the lowest swelling observed for GelMANB/GelSH is consistent with the higher storage modulus of this formulation and supports the formation of a more densely crosslinked or more effectively reinforced hydrogel network. Overall, these results show that the swelling behaviour of gelatin-based hydrogels can be tuned through the introduced functional groups and crosslinker chemistry, with GelMANB-based thiol-ene networks displaying reduced water uptake compared to GelMANB without additional thiolated crosslinker while maintaining gel fractions indicative of stable network formation.

#### 3.1.5 Tensile properties of crosslinked hydrogel networks

Tensile testing was performed to evaluate the macroscopic mechanical integrity and extensibility of the selected 10 w/v% hydrogel films under hydrated conditions. Rather than targeting a single tissue-specific tensile value, these measurements were used to compare the relative ability of the different gelatin-based networks to withstand deformation during handling, culture and potential application in mechanically dynamic soft tissue environments, such as lung-or urethra-inspired tissue engineering. The tensile properties of the 10 w/v% equilibrium-swollen hydrogel films were evaluated using ring-shaped samples.[36] This geometry was selected instead of conventional dogbone-shaped specimens to avoid clamping-induced premature failure, which is commonly encountered for soft gelatin-based hydrogels. Ultimate tensile strength, Young’s modulus and maximum strain at break were extracted from the resulting stress-strain curves (Figure 7).

**Figure 7.**
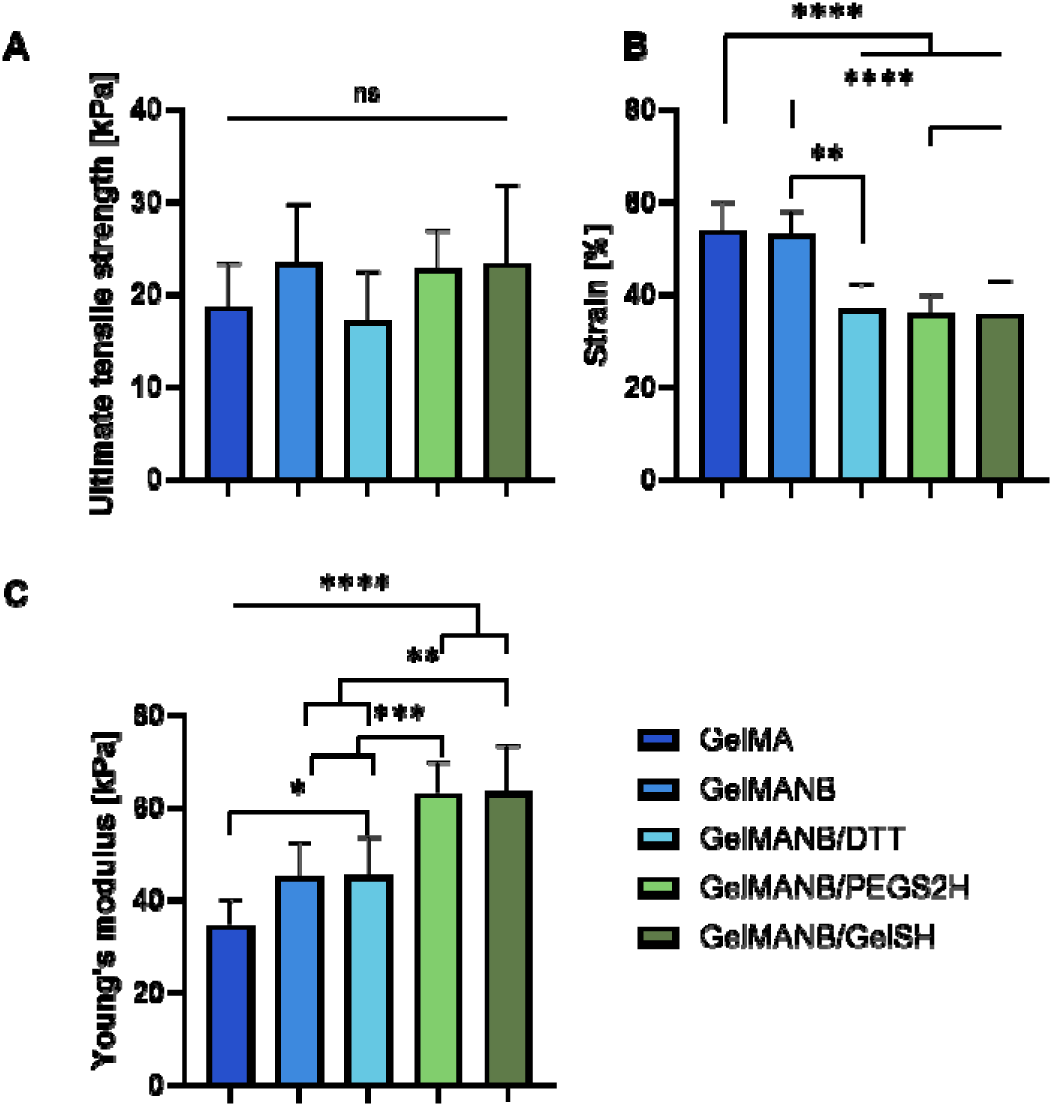
Tensile properties of 10 w/v% equilibrium-swollen GelMA and GelMANB-based hydrogel films. (A) Ultimate tensile strength, (B) maximum strain at break and (C) Young’s modulus were determined from stress-strain curves obtained from tensile testing on ring-shaped hydrogel samples. *Statistical analysis was performed using one-way ANOVA with Tukey’s post hoc test. *p < 0.05, **p < 0.01, ***p < 0.001 and ****p < 0.0001; ns indicates no significant difference*.

Overall, only limited differences in ultimate tensile strength were observed between the different hydrogel formulations, and no statistically significant differences were detected. This indicates that, under the applied testing conditions, the different crosslinking chemistries did not markedly alter the maximum stress that the hydrogels could withstand before failure. Therefore, crosslinker chemistry mainly influenced stiffness and extensibility rather than ultimate tensile strength.

In contrast, the Young’s modulus was clearly influenced by crosslinking chemistry. GelMANB/PEG2SH and GelMANB/GelSH showed comparable Young’s moduli of 62.98±6.89 kPa and 63.39±9.98 kPa, respectively. These values were higher than those obtained for GelMANB/DTT (45.21±8.39 kPa) and GelMANB without thiolated crosslinker (45.01±7.35 kPa). This may be related to the small bifunctional structure of DTT, which can form covalent thiol-ene crosslinks but does not introduce a polymeric spacer or macromolecular reinforcing phase. In contrast, PEG2SH provides a flexible polymeric spacer, while GelSH introduces a multifunctional gelatin-based crosslinker, both of which can contribute more effectively to tensile stiffening. This trend is consistent with the rheological data, where PEG2SH and GelSH also resulted in more reinforced networks.

The maximum strain at break followed a different trend. GelMA and GelMANB without thiolated crosslinker showed the highest extensibility, with strain values of 53.86±6.02% and 53.02±5.11%, respectively. In contrast, the thiol-ene-crosslinked GelMANB networks showed lower strain at break, indicating reduced extensibility after incorporation of additional thiol-ene crosslinks. This is consistent with the increase in Young’s modulus observed for the PEG2SH- and GelSH-crosslinked networks and suggests that additional covalent crosslinking restricted network deformability. Although chain-growth GelMA networks are often described as more heterogeneous than step-growth thiol-ene networks, this did not translate into lower extensibility in the present tensile tests. GelMA and GelMANB without thiolated crosslinker showed comparable strain at break, whereas the thiol-ene-crosslinked GelMANB formulations displayed reduced extensibility, suggesting that the additional covalent thiol-ene crosslinks increased network restriction and limited deformation before failure.[50]

Together, these tensile data confirm that the mechanical properties of GelMANB-based hydrogels can be tuned by varying the thiolated crosslinker. While ultimate tensile strength remained largely comparable across formulations, PEG2SH and GelSH increased the Young’s modulus, whereas the presence of thiol-ene crosslinks reduced the maximum strain at break. Importantly, GelMANB/DTT and GelMANB/GelSH displayed distinct mechanical profiles and network architectures, despite both remaining within a soft hydrogel range relevant for soft tissue engineering. On this basis, these two formulations were selected for subsequent digital light processing and biological evaluation to investigate how differences in network architecture and crosslinker chemistry influence printability and fibroblast response.

### 3.2 Optimization of resin compositions and DLP printing parameters

The processability of the selected resin formulations was evaluated using 10 w/v% GelMA, 10 w/v% GelMANB/DTT, 10 w/v% GelMANB/GelSH and 7.5 w/v% GelMANB/GelSH. These formulations were selected based on their physicochemical properties and their relevance for generating DLP-printed scaffolds within the mechanical range targeted for soft tissue engineering applications, including lung- and urethra-inspired models. DTT and GelSH were selected as thiolated crosslinkers for GelMANB to compare a small bifunctional crosslinker with a gelatin-based macromolecular crosslinker. GelSH was included because its collagen-derived backbone contains cell-interactive motifs, including RGD sequences that support cell adhesion, and because it avoids potential phase separation associated with combining gelatin and PEG-based crosslinkers. For all selected formulations, LAP and tartrazine concentrations were defined based on previous gelatin-based DLP literature [39–41,51,52] and formulation-specific photoabsorber screening, as described in Section 2.4.1 and summarised in Table 1. Briefly, 10 mol% LAP was used for all formulations, while tartrazine was selected at 2.0 mol% for GelMA-based resins and 2.44 mol% for GelMANB-based resins. These conditions were then used to evaluate working curves and scaffold printability.

Exposure time per layer is one of the key parameters that directly can influence the preciseness of the CAD/CAM design reproduction by the printed construct while also affecting cell survival. Thus, we consequently aimed to identify the shortest exposure time that still produced reliable layers and good CAD/CAM mimicry, as longer exposures and the overall longer printing time can reduce viability of the encapsulated cells in subsequent experiments.[53] Exposure time per layer was optimized through a two-stage approach. First, working curves were established for each selected resin to determine the penetration depth of the curing light (D_p_) and the minimal time required for crosslinking (t_c_), as shown in Figure S4. The minimal time required for crosslinking varied between the tested resin formulations: 6.48 s for 10 w/v% GelMA, 12.61 s for 10 w/v% GelMANB/DTT, 3.21 s for 10 w/v% GelMANB/GelSH and 5.58 s for 7.5 w/v% GelMANB/GelSH. The fastest crosslinking was observed for 10 w/v% GelMANB/GelSH, which is consistent with the photorheology data showing the highest crosslinking rate for this formulation.

Next, printability was further evaluated using more complex test structures, including the benchmark design and tissue engineering scaffolds shown in Figure 2C,D. These structures were printed at the minimal curing time determined from the working curves and at an increased exposure time of 14 s, since a slightly longer exposure can improve the printing reliability of more complex geometries.[54] Figure 8 shows the benchmark structures containing round and square pores printed from the different resin formulations at their minimal curing time and at 14 s exposure time. The obtained curing and exposure times are in the same order of magnitude as previously reported for gelatin-based DLP resins, where exposure conditions strongly depend on resin composition, photoinitiator concentration, photoabsorber use, light intensity and layer thickness.[51,52,55] For example, reported exposure times for 10 w/v% GelMA-based resins range from approximately 11 s to 30 s or longer, while Maes et al. [39] reported DLP processing of 10 w/v% GelMANB using an exposure time of 10.5 s at a light intensity of 36.4 mW·cm⁻². Based on the working curves and benchmark printing results, an exposure time of 14 s per layer at 37.6 mW·cm⁻² was selected for all four resin formulations to ensure reliable printing of the scaffold geometry while allowing direct comparison between formulations.

**Figure 8.**
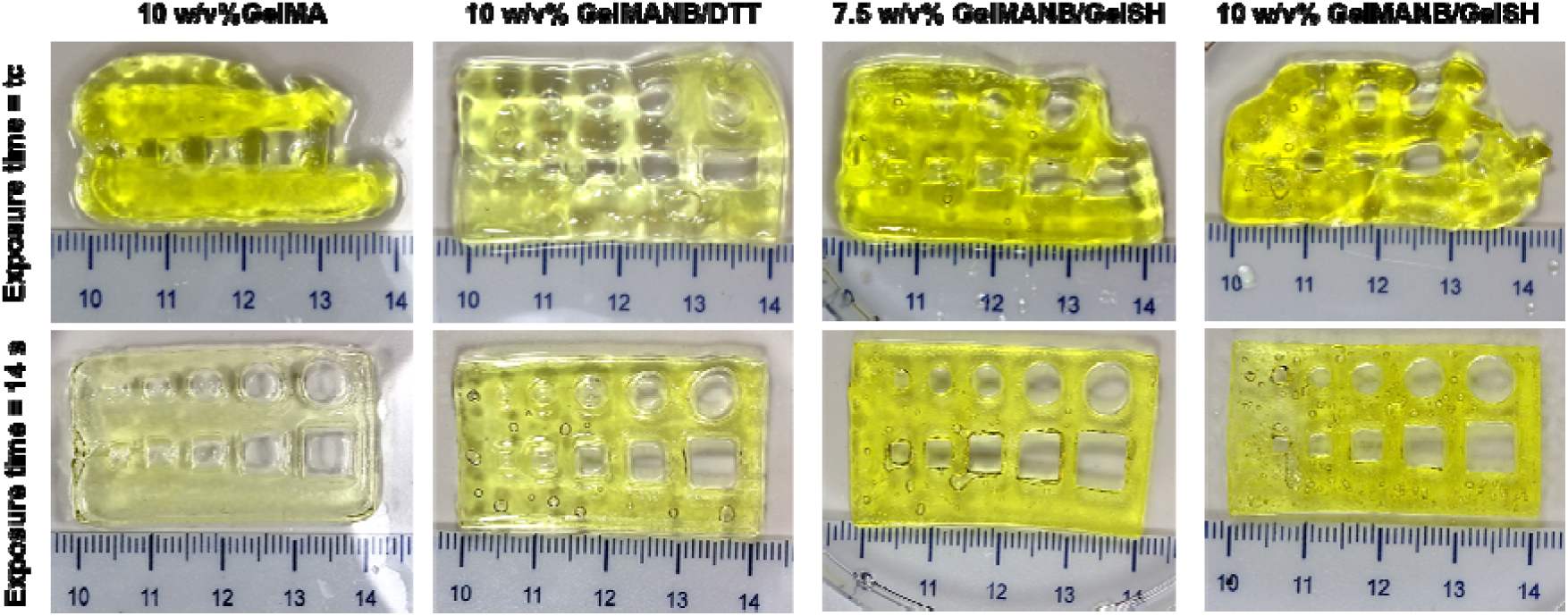
Benchmark design DLP printed using both critical (t_c_) and optimized (14 s) exposure times.

Constructs printed at critical energy dose for polymerisation exhibited severe mechanical fragility during handling and compromised edge definition, whereas those printed at 14 s exposure demonstrated superior structural integrity and feature sharpness. Moreover, all pores in the benchmarks printed at the higher exposure time remained visible and open, except for the smallest features (a 1 mm diameter circular pore and a square pore with a 1 mm side length). In contrast, at the minimal exposure time, the pores were markedly deformed and partially occluded by polymerised hydrogel. This improved feature fidelity at higher exposure may be attributed to increased crosslinking density, resulting in a more dimensionally stable network that better preserved the designed geometry without inducing the loss of resolution typically associated with overcuring.

Next, we evaluated the influence of the exposure time on the printability of TE scaffolds (CAD-Figure 2C). Equally as for the benchmark structure, constructs were manufactured using minimal exposure time and an exposure time at 14 s. In all resins tested, printing with minimal exposure time failed to reproduce the complex structure of TE scaffold. In contrast, longer exposure times produced constructs with higher resolution (Figure 9). As higher exposure time resulted in superior printing accuracy for both benchmark and TE scaffold, it was selected for further experiments.

**Figure 9.**
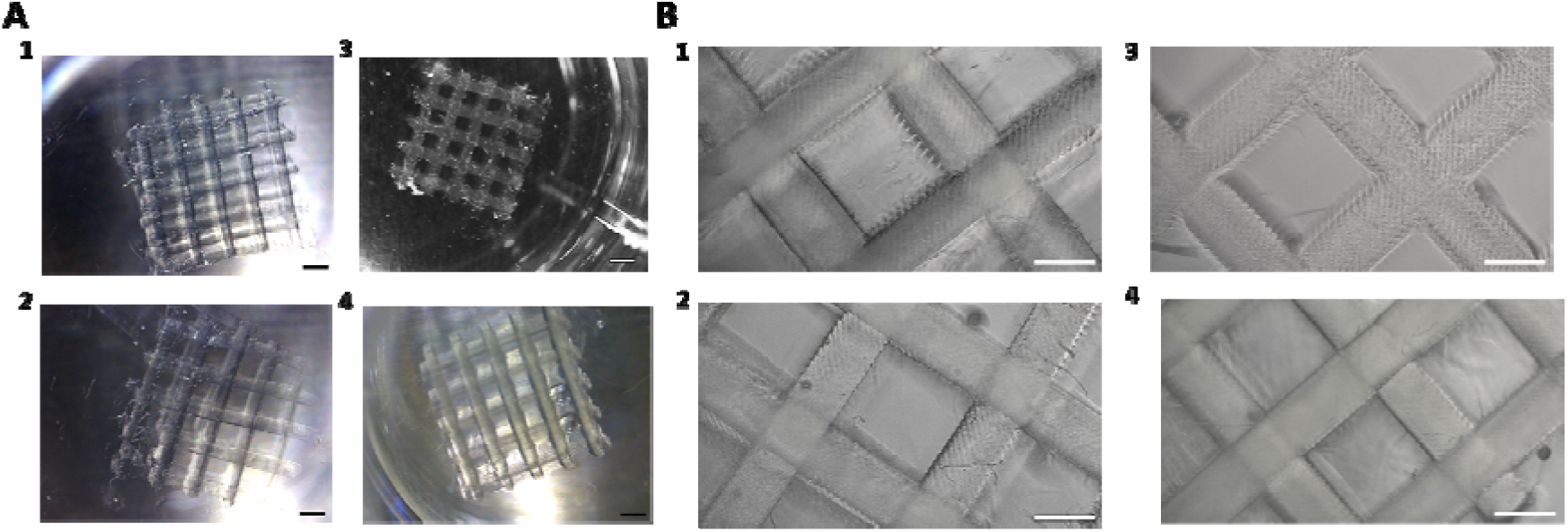
Light microscopy images of the DLP-printed tissue engineered constructs after washing steps: 1) 10 w/v% GelMA; 2) 10 w/v% GelMANB/DTT; 3) 7.5 w/v% GelMANB/GelSH; 4) 10 w/v% GelMANB/GelSH. (A) scale bar 1000 µm, (B) scale bar 500 µm.

Next, to assess the discrepancy between the CAD design and the DLP-printed tissue engineering scaffolds, samples were manufactured from the four selected resin formulations (Table 1) using the optimised printing parameters, including an exposure time of 14 s and a light intensity of 37.6 mW·cm⁻². The exposure time of 14 s was selected based on the working curve analysis and benchmark printing experiments described above, as the minimal curing times were sufficient for layer formation but did not provide reliable reproduction of the more complex scaffold architecture. Since the different formulations displayed distinct swelling behaviour, CAD/CAM mimicry was evaluated after equilibrium swelling in 1×, 3× and 10× PBS by measuring both strut and pore dimensions using optical microscopy (Figure 10).

**Figure 10.**
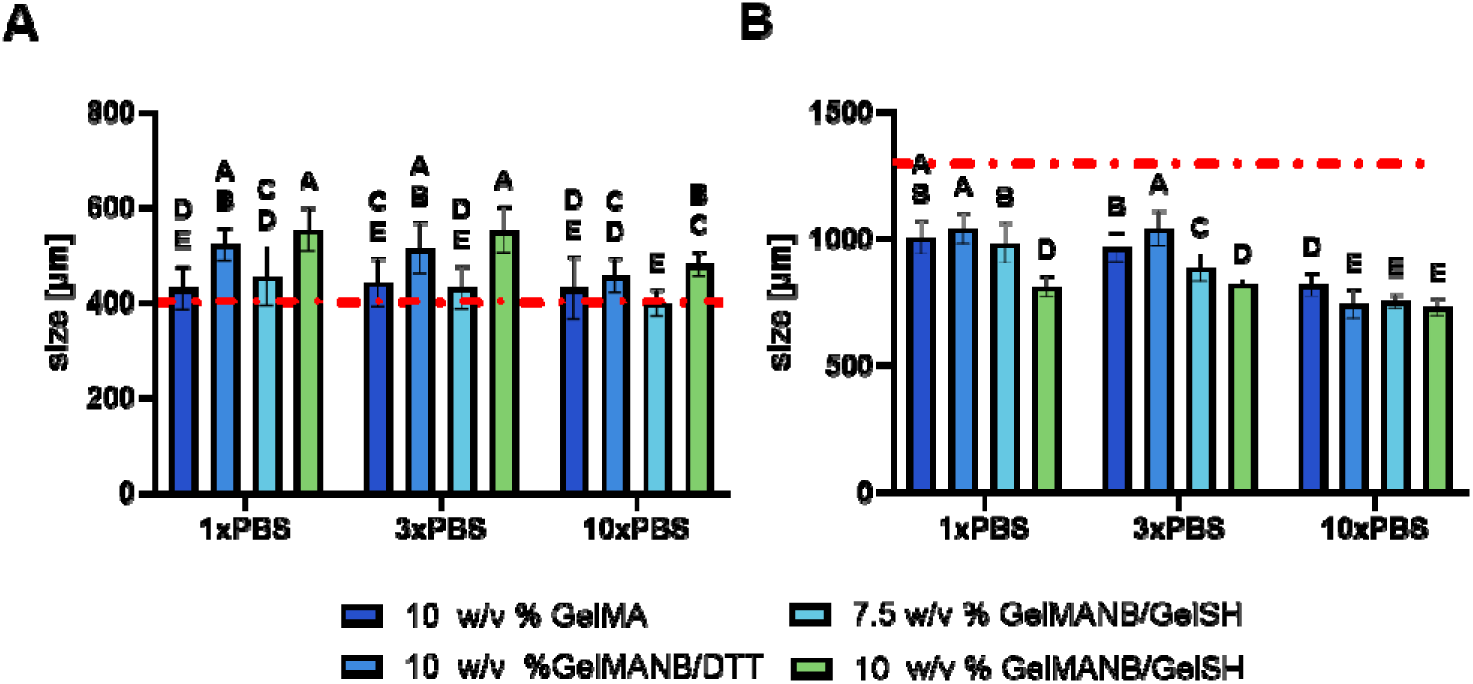
Measured strut (A) and pore (B) sizes of DLP-printed TE scaffolds equilibrated in PBS (1x, 3x, and 10x) via optical microscopy. Statistical analysis was performed using two-way ANOVA with Bonferroni’s multiple comparisons test. The horizontal red line indicates the (theoretical) pore or strut size of the CAD design. Different letters indicate significant differences at p < 0.05.

DLP-printed scaffolds prepared from 10 w/v% GelMA and 7.5 w/v% GelMANB/GelSH showed the closest agreement with the CAD strut dimensions, reaching 107.42% and 113.57% of the designed strut size, respectively. These values are within the ±15% tolerance commonly considered acceptable for hydrated polymeric scaffolds.[56] In contrast, scaffolds printed from 10 w/v% GelMANB/DTT and 10 w/v% GelMANB/GelSH showed larger deviations from the CAD design, with strut dimensions of 130.62% and 138.27%, respectively. These results indicate that, compared with GelMA, the 10 w/v% GelMANB/DTT and 10 w/v% GelMANB/GelSH formulations showed less accurate strut-size mimicry after swelling.

All DLP-printed tissue engineering scaffolds showed lower pore dimensions compared with the CAD design, which is consistent with the observed increase in strut size after swelling. The pore dimensions of scaffolds printed from 10 w/v% GelMA, 10 w/v% GelMANB/DTT and 7.5 w/v% GelMANB/GelSH were comparable, reaching 77.38%, 80.07% and 75.57% of the designed pore size, respectively. In contrast, 10 w/v% GelMANB/GelSH showed the largest reduction in pore size, reaching only 62.37% of the CAD value. This indicates that 10 w/v% GelMANB/GelSH deviated most strongly from the intended pore geometry, whereas GelMA, GelMANB/DTT and 7.5 w/v% GelMANB/GelSH showed comparable pore-size mimicry. The larger strut dimensions observed for 10 w/v% GelMANB/DTT and 10 w/v% GelMANB/GelSH do not necessarily contradict the swelling behaviour observed for bulk hydrogel films, since CAD/CAM mimicry of printed scaffolds is not governed by swelling alone. In printed porous structures, the final dimensions also depend on resin-specific curing behaviour, local light propagation, overcuring around small features and the geometry of the scaffold itself. The improved CAD/CAM mimicry of 7.5 w/v% GelMANB/GelSH compared with 10 w/v% GelMANB/GelSH therefore suggests that lowering the total gelatin concentration reduced excessive strut enlargement under the selected printing conditions, resulting in a closer match with the intended CAD geometry.

Overall, these findings demonstrate that scaffold fidelity after swelling is formulation-dependent and is influenced by the combined effects of resin composition, curing behaviour, scaffold geometry and hydrogel swelling. Consequently, CAD designs may need to be adjusted for each resin formulation to compensate for swelling- and printing-induced changes in strut and pore dimensions and to achieve the desired final scaffold architecture.

The two intended application contexts, lung and urethral tissue engineering, are associated with markedly different fluid milieus. Lung tissue is exposed to near-physiological osmolarity, whereas urine osmolarity can vary widely and may reach hyperosmotic values in concentrated urine.[44] Therefore, dimensional stability was evaluated after incubation in 1×PBS (280-300 mOsm·L⁻¹) as a physiologically relevant condition for lung-inspired soft tissue engineering, 3×PBS (840-900 mOsm·L⁻ ¹) as a defined hyperosmotic condition within the range of concentrated human urine, and 10×PBS (2800-3000 mOsm·L⁻ ¹) as an accelerated stress condition to evaluate the robustness of the printed hydrogel networks beyond the physiological urine range. As osmotic conditions can affect the hydration, mesh size and mechanical stability of collagen, gelatin and gelatin derivatives such as GelMA [57], these conditions allowed us to compare scaffold dimensional stability across both physiological and hyperosmotic environments. At 1×PBS, all scaffolds retained their porous architecture, supporting dimensional stability under lung-relevant physiological osmolarity. At 3×PBS, no significant changes in strut or pore size were observed for most formulations, except for 7.5 w/v% GelMANB/GelSH, where pore size decreased from 75.57 to 68.09% of the initial CAD value. Under the non-physiological 10×PBS stress condition, all scaffolds exhibited a significant reduction in pore size, whereas changes in strut size were less pronounced. Overall, these findings indicate that the scaffolds largely retained dimensional stability under both lung-relevant physiological osmolarity and urethra-relevant hyperosmotic conditions, while 10×PBS revealed the expected osmotic shrinkage of the hydrogel networks. In this study, PBS-based media provided defined and reproducible osmotic environments, allowing dimensional changes to be primarily attributed to osmolarity rather than to the complex biochemical composition of tissue fluids or urine. Nevertheless, PBS does not fully reproduce either the lung or urethral environment. Future studies should therefore evaluate scaffold behaviour under more application-specific conditions, including lung-relevant culture media and artificial or donor urine [44], ideally using cell-laden scaffolds, as both medium composition and developing tissue may influence hydrogel stability and cellular response.

Next, the physico-chemical properties of scaffolds manufactured using the optimised printing conditions were evaluated. Cubic scaffolds (CAD geometry shown in Figure 2A) printed from all four resin formulations exhibited high gel fractions (>70%)(Figure 11A), indicating efficient formation of the crosslinked polymer networks. The mass swelling ratios were comparable for scaffolds based on 10 w/v% GelMA (20.83±0.83), 7.5 w/v% GelMANB/GelSH (19.34±0.90), and 10 w/v% GelMANB/GelSH (19.13±0.62), with no significant differences observed between these formulations. In contrast, GelMANB/DTT scaffolds exhibited a significantly higher mass swelling ratio (26.15±1.86) (Figure 11B), indicating greater water uptake by this network.

**Figure 11.**
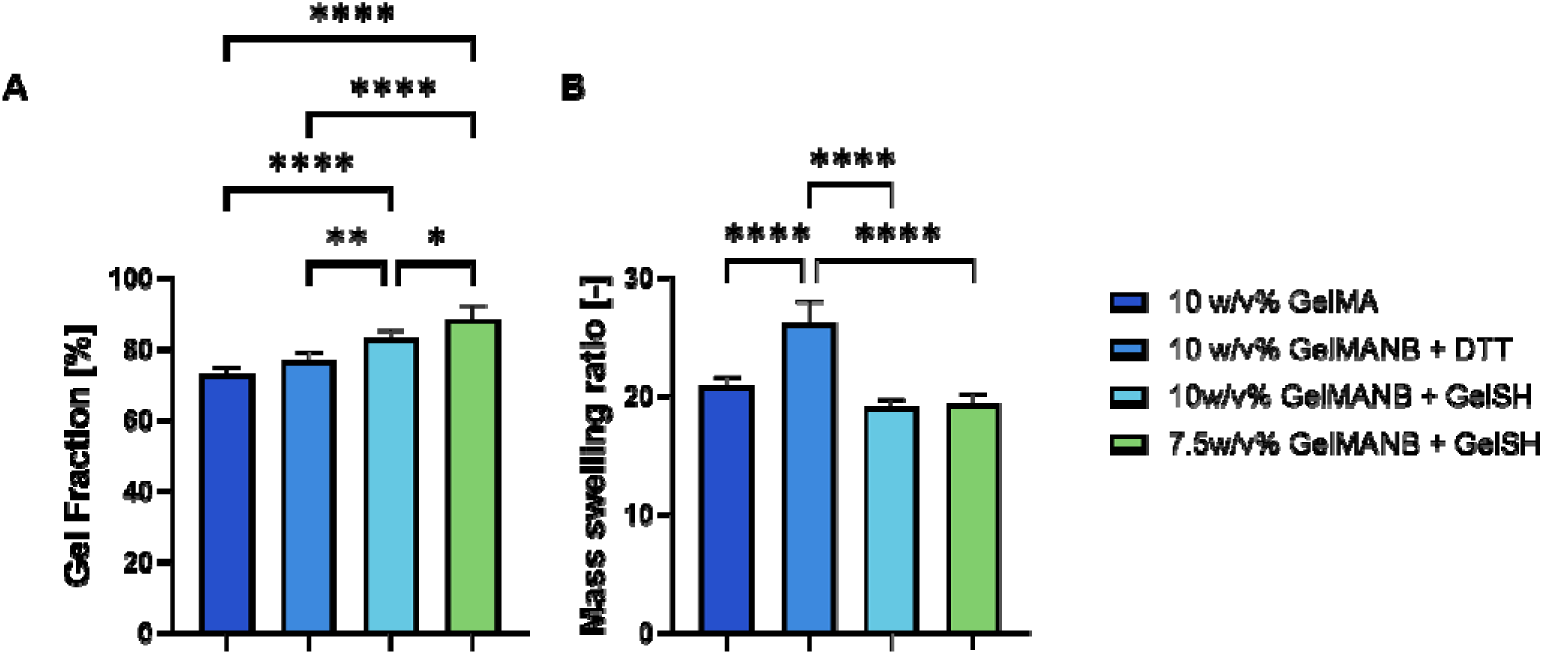
Gel fraction (A) and mass swelling ratio (B) of GelMA and GelMANB-based DLP-printed scaffolds. Statistical analysis was performed using two-way ANOVA with Tukey’s post hoc analysis: *p *< 0.05, **p < 0.01 and ****p < 0.0001*.

The different swelling behaviour of 10 w/v% GelMANB/DTT and 7.5 w/v% GelMANB/GelSH, despite their comparable G′ values, highlights that scaffold hydration is influenced not only by bulk stiffness, but also by crosslinker architecture and the resulting network structure. DTT is a small bifunctional crosslinker that can form covalent bridges between GelMANB chains, whereas GelSH acts as a gelatin-based macromolecular crosslinker and contributes additional polymeric network structure. This difference may explain why GelMANB/DTT scaffolds showed higher water uptake, while GelMANB/GelSH scaffolds displayed lower swelling at comparable or higher stiffness.

### 3.3. DLP bioprinting using human foreskin fibroblast (HFF-1)

#### 3.3.1. Effect of HFF-1 encapsulation in DLP-printed GelMA- and GelMANB-based scaffolds on the metabolic activity of the cells

For the *in vitro* assessment of the cytocompatibility of the developed resin compositions, human foreskin fibroblasts (HFF-1) were selected as a standardized stromal cell model to evaluate cell viability, morphology, and proliferative response following encapsulation and 3D bioprinting. The disc scaffolds (CAD model - Figure 2B) were manufactured using previously optimised resin formulations and printing conditions, presented in Table 2. Metabolic activity of HFF-1 cells, as an indicator of cell viability within DLP-printed scaffolds, was evaluated on days 2, 5, and 7 post-printing using MTS assay.

**Table 2.** Overview of the resin formulations and printing parameters used for the cell-laden bioprinting.

| Gelatin type | Total gelatin conc. [w/v%] | LAP conc. [mol%] | Tartrazine conc. [mol%] | TCEP [mol%] | Intensity [mW·cm<sup>-2</sup>] | Curing time /layer [s] | Printing bed T [°C] | HFF-1 final concentration [$\cdot 10^6$ cells·mL<sup>-1</sup> of resin] |
| --- | --- | --- | --- | --- | --- | --- | --- | --- |
| GelMA | 10 | 10 | 2 | ----- | 37.6 | 14 | 45 | 2 |
| GelMANB/DTT | 10 | 10 | 2.44 | 25 | 37.6 | 14 | 45 | 2 |
| GelMANB/GelSH | 7.5 | 10 | 2.44 | 25 | 37.6 | 14 | 45 | 2 |

On day 2 post-printing, no significant differences in metabolic activity were observed between cells encapsulated in the different resin formulations (Figure 12). By day 5, metabolic activity decreased for all formulations, indicating a transient reduction in cellular metabolic response after printing. Similar non-linear trends in metabolic activity have been reported for other cell-laden bioprinted hydrogel systems, where an initial decrease or lag phase was attributed to cell adaptation to the printed 3D microenvironment, followed by partial recovery or increased metabolic activity at later time points.[58] In the present study, the day-5 decrease may reflect cellular adaptation to the post-printing microenvironment and/or exposure to residual resin components, such as photoinitiator [59], photoabsorber [60], or unreacted crosslinking components [29]. Although DTT-containing systems have previously raised concerns regarding potential cytotoxicity when residual or freely soluble DTT is present [29], the present data do not indicate a stronger reduction in metabolic activity for GelMANB/DTT compared with the other formulations. Therefore, the observed day-5 decrease is more likely related to a combination of printing-associated stress, early cellular adaptation and formulation-dependent mass transport rather than to one specific resin component. Limitations in oxygen and nutrient transport within the 3D scaffolds may additionally contribute to reduced cellular metabolic activity.[61] By day 7, metabolic activity increased again in all formulations, reaching levels comparable to those observed on day 2. As metabolic activity alone does not directly reflect cell number or viability and can also be influenced by cellular stress [62], cell viability and morphology were additionally assessed by Calcein AM/PI staining on days 2, 5, and 7 (Figure 13).

**Figure 12.**
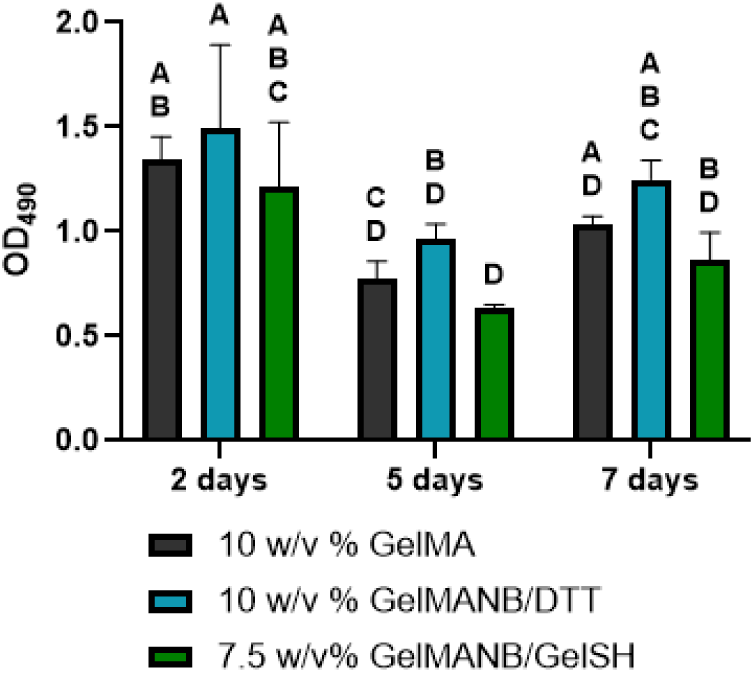
Metabolic activity of HFF-1 encapsulated in DLP-printed GelMA- and GelMANB–based scaffolds at day 2, day 5, and day 7 post-encapsulation. Statistical analysis was performed using two-way ANOVA with Tukey’s post-hoc test. Data are expressed as mean±SD (n = 3). Different letters indicate significant differences at p < 0.05.

**Figure 13.**
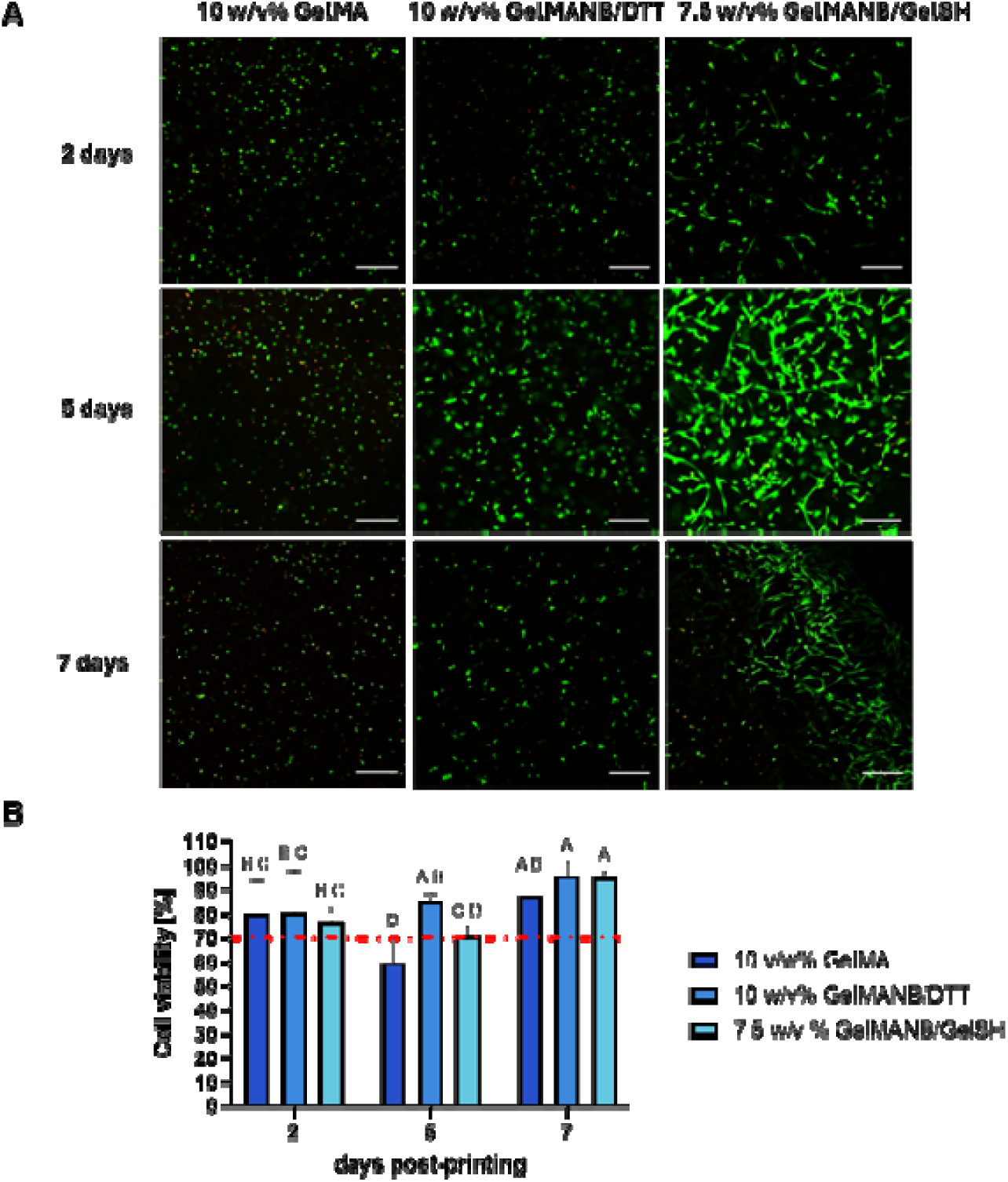
(A) CLSM images of HFF-1 encapsulated in scaffolds manufactured by DLP-printing from different resin formulations. Viability was evaluated using staining with calcein AM/PI on days 2,5, and 7 post-printing. Images are merged for viable (green) and dead (red) cells. Scale bar represents 200 μm. (B) Quantification of live and dead cells on day 2, 5, and 7 post-printing of the DLP-printed scaffolds (disks). According to the ISO standard (ISO 10993–5), viability above 70% is required (cut-off indicated on the graphs). Statistical analysis was performed using two-way ANOVA with Bonferroni’s multiple comparisons test. Data are expressed as mean±SD (n = 14). Different letters indicate significant differences at p < 0.05.

On day 2, the majority of HFF-1 cells remained viable in all formulations, with viability exceeding 70% (Figure 13B). This is consistent with the 70% viability threshold commonly applied in cytotoxicity assessment according to ISO 10993-5:2009. Notably, cells encapsulated in GelMANB/GelSH already showed initial spreading, with the appearance of elongated fibroblast morphology (Figure 13A).

By day 5, cell viability in the GelMA-based scaffolds decreased below 70%, consistent with the concomitant reduction in metabolic activity (Figure 14). In contrast, viability in both GelMANB-based formulations remained at or above 70%, with only minor changes compared with day 2. The transient decrease in metabolic activity of cells in the GelMANB-based scaffolds therefore occurred without a corresponding pronounced loss of membrane integrity. This may indicate temporary cellular adaptation to the 3D microenvironment and/or a reduced metabolic state associated with lower cellular stress, rather than extensive cell death. Differences in cell morphology were also apparent: cells in GelMA remained predominantly rounded, whereas spreading was observed in GelMANB/DTT, and the majority of cells in GelMANB/GelSH displayed the characteristic spindle-shaped morphology of fibroblasts.

**Figure 14.**
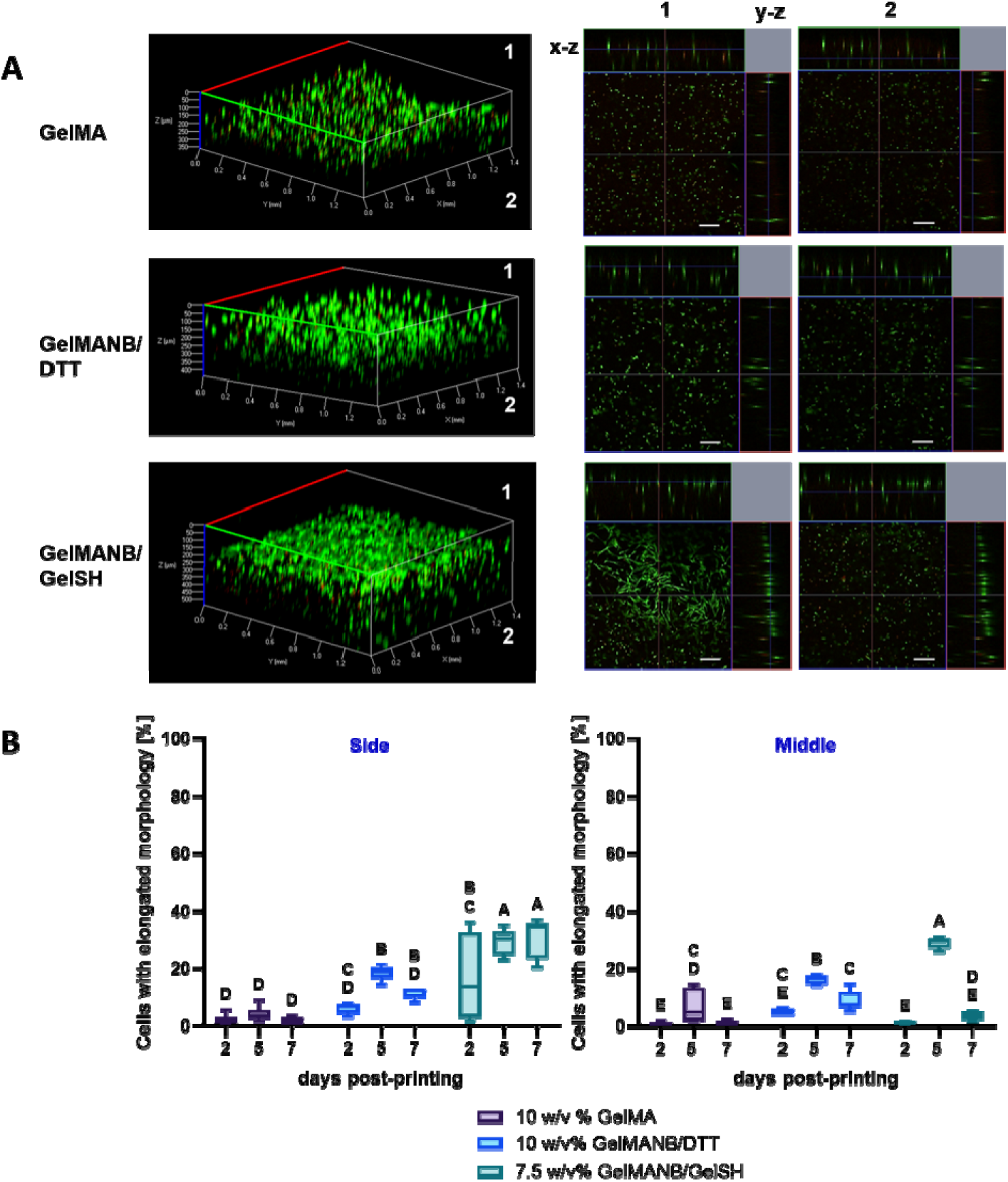
Distribution and morphology of the encapsulated HFF-1 cells on the 7-day post-printing. (A) Z-stack reconstitution and orthogonal (XZ,YZ) projections of calcein AM(green)/PI(red)-stained HFF-1 within DLP-printed scaffolds. Orthogonal views reveal depth-dependent differences in cell morphology at upper (1) and deeper (2) layers. Scale bars correspond to 200 µm. (B) Percentage of the cells displaying elongated morphology after encapsulation in 10 w/v% GelMA, 10w/v% GelMANB/DTT and 7.5 w/v% GelMANB/GelSH analyzed on two different layers of the printed scaffold (in outer layers (side) and within the interior (middle)). Measurements were taken on day 2,5 and 7 post-printing (n=6). Statistical analysis was performed using two-way ANOVA with Bonferroni’s multiple comparisons test. Different letters indicate significant differences at p < 0.05.

By day 7, high cell viability was observed in both GelMANB-based formulations, reaching 95.0±5.9% for GelMANB/DTT and 94.9±2.7% for GelMANB/GelSH. Cells encapsulated in GelMA exhibited a slightly lower viability of 87.2±6.0%, although this difference was not significant. Overall, the recovery in metabolic activity together with the high viability observed at day 7 demonstrates that HFF-1 cells remained compatible with all investigated formulations over the evaluated culture period.

Although no substantial differences in cell viability were observed between the two GelMANB crosslinking systems, more pronounced cell spreading was observed when GelSH was used as the crosslinker. This may partly result from the additional cell-adhesive motifs introduced by GelSH, including arginine-glycine-aspartic acid (RGD) sequences that promote integrin-mediated cell adhesion and spreading [45]. In addition, the step-growth thiol-ene crosslinking mechanism of GelMANB-based networks produces a more homogeneous network architecture than the chain-growth polymerisation of GelMA and is associated with lower radical concentrations during network formation. Chain-growth GelMA polymerisation can generate heterogeneous kinetic chains that may restrict cellular access to enzymatically cleavable sites within the gelatin backbone [63], while the higher radical burden may additionally contribute to cellular stress during encapsulation and printing, potentially limiting cell-mediated matrix remodelling and spreading.

Further analysis of reconstructed CLSM Z-stacks revealed pronounced differences in cell morphology across the scaffold depth (Figure 14). As HFF-1 fibroblasts typically adopt an elongated, spindle-shaped morphology, cell circularity was used to quantify spreading, with cells exhibiting a circularity <0.5 classified as elongated (Figure 14B). In GelMA-based scaffolds, cells remained predominantly rounded throughout the 7-day culture period, with only a transient increase in elongated cells within the inner regions on day 5 (day 2: 0.59±0.50%; day 5: 6.61±5.20%).

In the thiol-ene-crosslinked scaffolds, cell morphology changed more markedly over time. In GelMANB/DTT scaffolds, the proportion of elongated cells increased from 4.9±1.0% on day 2 to 16.2±1.5% on day 5, with spreading observed both near the scaffold surface and within the inner regions. By day 7, this proportion decreased to 8.6±3.2%, which was not significantly different from day 2. GelMANB/GelSH scaffolds showed the most sustained cell spreading, with the proportion of elongated cells increasing from 16.5±13.2% on day 2 to 29.4±4.1% on day 5 and remaining elevated at 30.9±6.3% on day 7.

Importantly, cell morphology in the GelMANB scaffolds was strongly dependent on the position within the 2-mm-thick samples. Although the total gelatin concentration was kept constant at 7.5 w/v%, this formulation consisted of both GelMANB and GelSH, whereas GelMANB/DTT contained GelMANB as the only gelatin-based polymer component. The pronounced fibroblast elongation observed in the GelMANB/GelSH system may therefore result from a combination of factors, including the presence of GelSH-derived cell-adhesive motifs, the gelatin-based macromolecular crosslinker architecture and differences in network organisation that may support cell-mediated matrix remodelling. However, elongated cells were predominantly located near the outer surfaces, whereas cells in the inner regions remained largely rounded despite exhibiting comparable viability. Day-7 Z-stack reconstructions confirmed this surface-to-core transition, with elongated cells mainly confined to approximately the outermost 150-200 µm and rounded cells predominating at greater depths (Figure 14A). These findings indicate that material chemistry alone is insufficient to support uniform fibroblast elongation throughout thicker hydrogel structures and suggest that limitations in oxygen and/or nutrient transport may contribute to the observed depth-dependent morphology.

To investigate whether reducing diffusion distances could improve cell spreading, a porous cubic scaffold with a strut size of 400 µm was subsequently evaluated (Figure 2C), substantially reducing the characteristic hydrogel thickness compared with the 2-mm-thick discs. Maximum-intensity projections of CLSM Z-stacks showed abundant elongated HFF-1 cells throughout the porous GelMA- and GelMANB-based scaffolds (Figure 15A). Morphometric analysis further demonstrated that elongated cells constituted the majority of the cell population. Compared to GelMA-based scaffolds, GelMANB/GelSH-based scaffolds showed a significantly higher number of cells with elongated morphology (Figure 15B).

**Figure 15.**
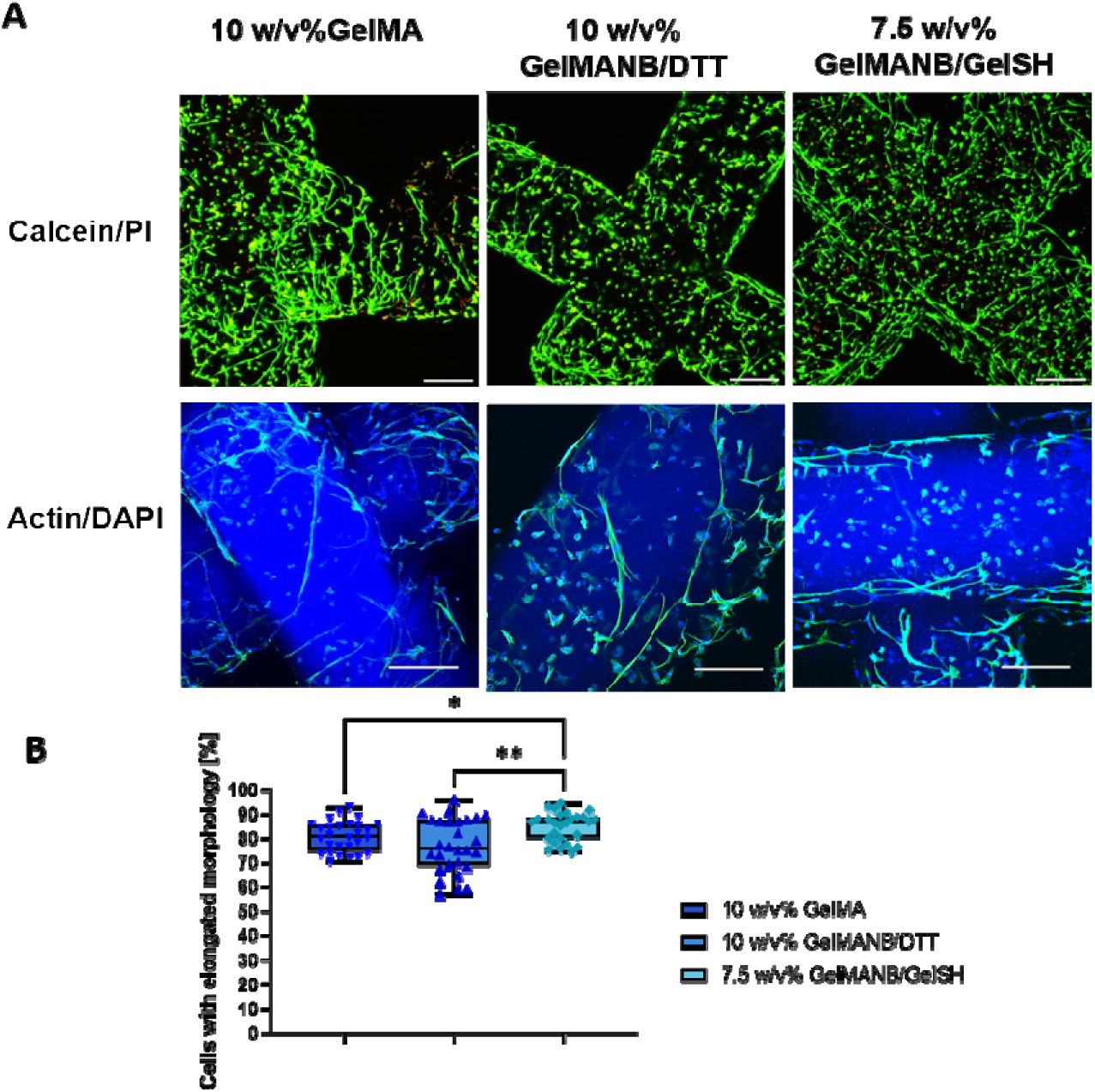
(A) Maximum intensity projections (MIP) of CLSM image stacks acquired to characterise colonization of the DLP-printed scaffolds by HFF-1 on day 7 post-printing. Colour allocation: Calcein/PI: green - living cells, red - dead cells; Actin/DAPI: green - actin, blue - nucleus. Scale bar- 200 µm. (B) Percentage of the cells displaying elongated morphology after encapsulation in 10 w/v% GelMA, 10 w/v% GelMANB/DTT, and 7.5 w/v% GelMANB/GelSH. Measurements were taken on day 7 post-printing (n=28). Statistical analysis was performed using Brown-Forsythe and Welch’s ANOVA, followed by Dunnett’s T3 multiple comparisons test. * p < 0.05, **- p < 0.005

Together, these results indicate that both hydrogel chemistry and scaffold architecture influence HFF-1 morphology. While GelMANB/GelSH promoted greater cell spreading in the thicker disc geometry, this advantage was strongly depth-dependent. Reducing the diffusion distance through a thinner, porous architecture markedly increased the proportion of elongated cells across all formulations. These findings highlight scaffold geometry as an important design parameter for supporting homogeneous cellular behaviour throughout 3D-printed tissue-engineered scaffolds.

## 4. Conclusion

In this work, bifunctional methacryloyl-norbornene gelatin (GelMANB) was developed as a modular photocrosslinkable platform for cell-laden digital light processing (DLP) of soft tissue-engineered scaffolds. Polymer concentration and thiolated crosslinker selection enabled substantial tuning of the network properties, with storage moduli ranging from 0.23 kPa for 5 w/v% GelMANB to 16.23 kPa for 10 w/v% GelMANB/GelSH. Gelatin methacryloyl (GelMA, 96% methacryloyl substitution) was used as a benchmark for GelMANB (68% methacryloyl and 28% norbornene substitution), providing comparable overall degrees of functionalisation. GelMANB/DTT at 10 w/v% (G′ = 8.63 kPa) and GelMANB/GelSH at 7.5 w/v% (G′ = 8.06 kPa) were selected for DLP processing and cell-laden printing based on their relatively comparable storage moduli to 10 w/v% GelMA (G′ = 11.85 kPa), while 10 w/v% GelMANB/GelSH (G′ = 16.23 kPa) was included to evaluate the influence of polymer concentration.

Optimised DLP conditions enabled reproducible fabrication of porous scaffolds, with strut dimensions reaching 107.42% and 113.57% of the CAD dimensions for 10 w/v% GelMA and 7.5 w/v% GelMANB/GelSH, respectively. These results indicate that GelMA and 7.5 w/v% GelMANB/GelSH provided the closest strut-size mimicry among the tested formulations, whereas pore dimensions were more strongly affected by formulation-dependent swelling. The scaffolds largely retained their dimensions under 1×PBS, representing physiological osmolarity relevant to lung-inspired applications, and under 3×PBS, representing a defined hyperosmotic condition relevant to concentrated urine. This supports their dimensional stability across both lung- and urethra-relevant osmotic conditions, while the non-physiological 10×PBS condition served as an accelerated stress test to reveal osmotic shrinkage of the hydrogel networks.

All formulations supported HFF-1 encapsulation and DLP printing, with day-7 cell viability reaching 95.0±5.9% for GelMANB/DTT, 94.9±2.7% for GelMANB/GelSH and 87.2±6.0% for GelMA. Compared with GelMA, the GelMANB formulations therefore showed higher day-7 viability under the investigated conditions. GelMANB/GelSH promoted the most sustained fibroblast spreading in 2-mm-thick samples, with elongated cells increasing from 16.5±13.2% on day 2 to 30.9±6.3% on day 7. However, spreading was predominantly located near the outermost 150-200 µm. Reducing the diffusion distance using porous scaffolds with 400 µm struts markedly enhanced cell spreading across all formulations, highlighting the importance of scaffold architecture alongside material chemistry.

Overall, GelMANB provides a versatile gelatin-based platform in which polymer concentration and crosslinker chemistry can be used to tailor network properties while maintaining DLP processability and cytocompatibility. Compared with the GelMA benchmark, GelMANB did not uniformly improve CAD/CAM mimicry, but provided additional crosslinker-dependent control over hydrogel properties and supported improved fibroblast viability and elongation. These findings highlight the combined importance of gelatin chemistry, printing conditions and scaffold architecture when designing bioinks for cell-laden DLP of soft tissue engineering platforms.

## 5. Acknowedgements

N. Pien and S. Van Vlierberghe would like to acknowledge Horizon Europe for funding the STRONG-UR project (Grant agreement ID: 101191695). STRONG-UR is a project funded by the European Union and has received funding from the Horizon Europe programme under grant agreement No 101191695. Views and opinions expressed are however those of the author(s) only and do not necessarily reflect those of the European Union or HaDEA. Neither the European Union nor the granting authority can be held responsible for them. N.Pien would like to acknowledge the financial support of the Research Foundation Flanders (FWO) under the form of an FWO senior postdoctoral fellowship (12A1U27N). We thank the NMR expertise centre (Ghent University) for providing support and access to its NMR infrastructure. The 400 and 500 MHz used in this work have been funded by a grant of the Research Foundation Flanders (FWO) (grant number FWO I006920N (GISMO code 319309020) and G011015N), and the ‘Bijzonder Onderzoeksfonds’ (BOF) (grant number BOF.BAS.2022.0023.01 (GISMO code BOF/BAS/2022/105) and BOF.BAS.20200019.01 (GISMO code 01B07520)). We thank BioRender.com for providing the platform used to create the graphical abstract in this manuscript.

## 7. Supplementary information

**Figure S1.**
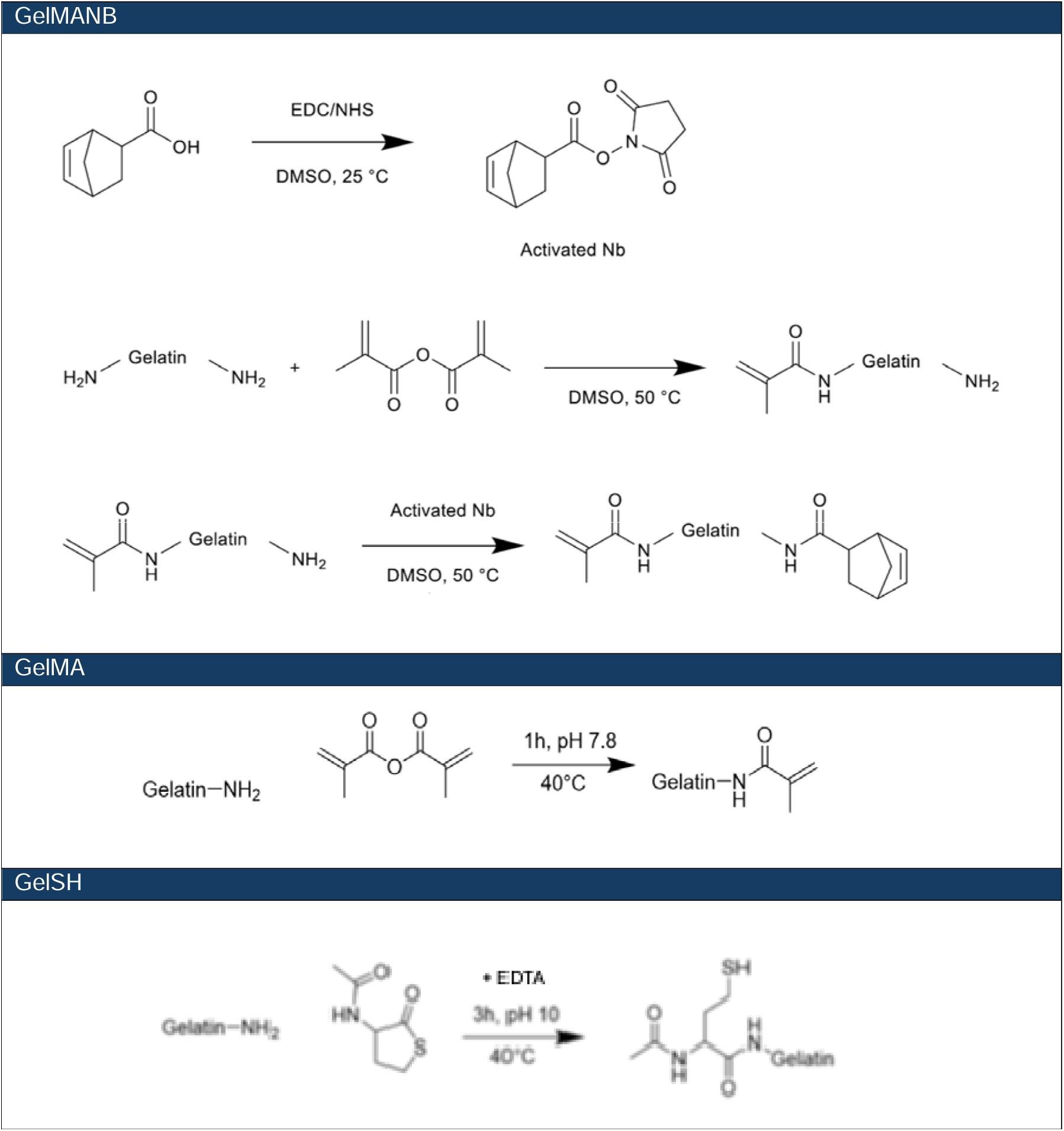
Reaction scheme showing the development of GelMANB, GelMA and GelSH; EDC: N-(3-dimethylaminopropyl)-N’-ethylcarbodiimide hydrochloride; NHS: N-hydroxysuccinimide, EDTA: ethylenediaminetetraacetic acid.

**Figure S2.**
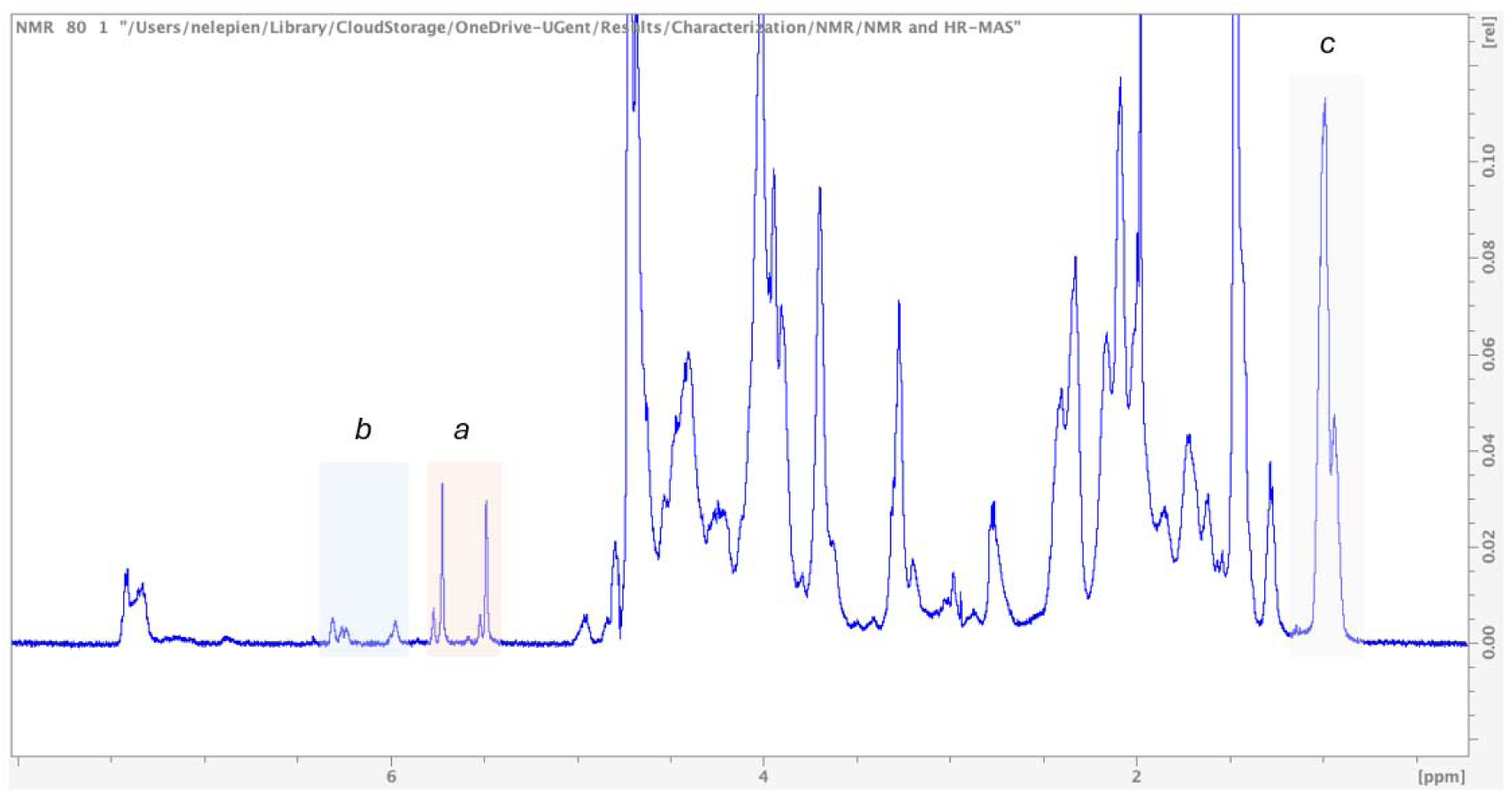
^1^H-NMR spectrum of GelMANB. The peaks corresponding to the protons of the methacryloyl functionalities are indicated with a and the norbornene functionalities are indicated with *b. The peak that is marked with c corresponds to the hydrogens from the chemically inert valine, leucine, and isoleucine amino acids*.

**Figure S3.**
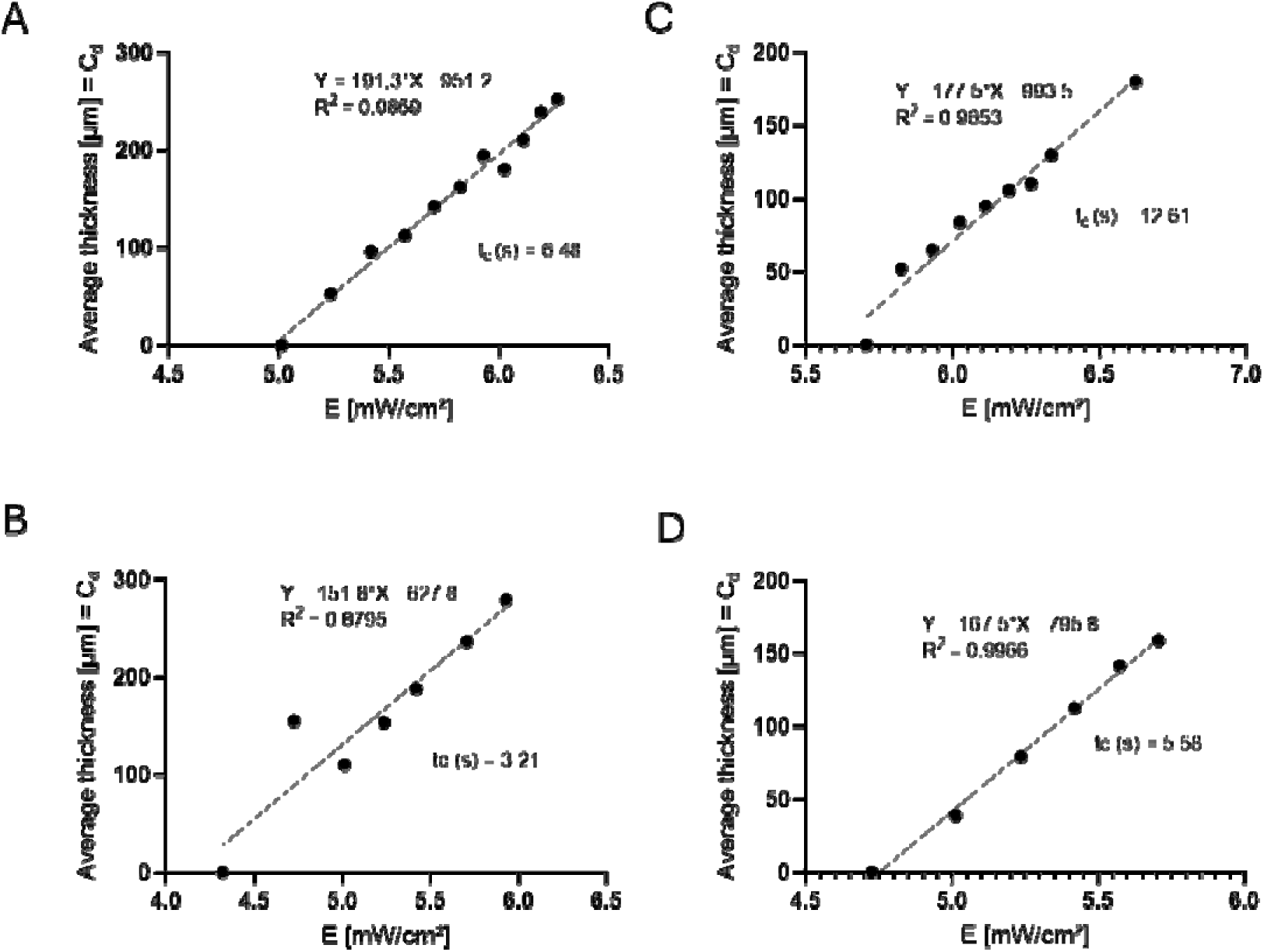
Working curves describing the relation between the applied dose (E) and the cured thickness (Cd) of the resins: (A) 10 w/v% GelMA, 10 mol% LAP, 2 mol% tartrazine; (B) 10 w/v% *GelMANB/GelSH, 10 mol% LAP, 2.44 mol% tartrazine; (C) 10 w/v% GelMANB/DTT, 10 mol% LAP, 2.44 mol% tartrazine; (D) 7.5 w/v% GelMANB/GelSH, 10 mol% LAP, 2.44 mol% tartrazine*.

## Notes

### Competing Interest Statement

The authors have declared no competing interest.

